# Explainable machine learning predictions to help anesthesiologists prevent hypoxemia during surgery

**DOI:** 10.1101/206540

**Authors:** Scott M. Lundberg, Bala Nair, Monica S. Vavilala, Mayumi Horibe, Michael J. Eisses, Trevor Adams, David E. Liston, Daniel King-Wai Low, Shu-Fang Newman, Jerry Kim, Su-In Lee

**Affiliations:** Paul G. Allen School of Computer Science and Engineering, University of Washington, Seattle, WA, USA.; Department of Anesthesiology and Pain Medicine, University of Washington, Seattle, WA, USA.; Seattle Children’s Hospital, Seattle, WA, USA.; Veterans Affairs Puget Sound Health Care System, Seattle, WA, USA.; Department of Genome Sciences University of Washington, Seattle, WA, USA.; Harborview Injury Prevention and Research Center, Seattle, WA, USA

## Abstract

Hypoxemia causes serious patient harm, and while anesthesiologists strive to avoid hypoxemia during surgery, anesthesiologists are not reliably able to predict which patients will have intraoperative hypoxemia. Using minute by minute EMR data from fifty thousand surgeries we developed and tested a machine learning based system called *Prescience* that predicts real-time hypoxemia risk and presents *an explanation of factors* contributing to that risk during general anesthesia. Prescience improved anesthesiologists’ performance when providing *interpretable hypoxemia risks with contributing factors*. The results suggest that if anesthesiologists currently anticipate 15% of events, then with Prescience assistance they could anticipate 30% of events or an estimated additional 2.4 million annually in the US, a large portion of which may be preventable because they are attributable to modifiable factors. The prediction explanations are broadly consistent with the literature and anesthesiologists’ prior knowledge. Prescience can also improve clinical understanding of hypoxemia risk during anesthesia by providing general insights into the exact changes in risk induced by certain patient or procedure characteristics. Making predictions of complex medical machine learning models (such as Prescience) interpretable has broad applicability to other data-driven prediction tasks in medicine.

## Introduction

Over 200 million surgeries are performed worldwide every year, with 30 million in the United States alone (*1*). Though an integral part of healthcare, surgery and anesthesia pose considerable risk of complications and death. Studies have shown a perioperative mortality rate of 0.4 to 0.8% and a complication rate of 3 to 17%, just in industrialized countries (*2*, *3*). Fortunately, half of these complications are preventable (*2*, *3*). With increasing adoption of electronic medical record systems, high fidelity heterogeneous data are being captured during surgery and anesthesia care, yet the utilization of these data to improve patient safety and quality of care remains poor (*4*). There is untapped potential for data science to utilize perioperative data to positively impact surgical and anesthesia care (*5*). Recognizing this unmet need and leveraging recent advances in perioperative informatics, we present new data science methods to predict harmful physiological events and so inform anesthesiologists.

Hypoxemia or low arterial blood oxygen tension is an unwanted physiological condition and can cause serious patient harm during general anesthesia and surgery (*6*). Hypoxemia is associated with cardiac arrest, cardiac arrhythmias, postoperative infections and wound healing impairments, decreased cognitive function and delirium, and cerebral ischemia through a number of metabolic pathways (*7*). Despite the advent and use of pulse oximetry to continuously monitor blood oxygen saturation (SpO_2_) during general and regional anesthesia, hypoxemia can neither be reliably predicted nor prevented (*8*). Real-time blood oxygen monitoring through pulse oximetry only allows anesthesiologists to take reactive actions to minimize the duration of hypoxemic episodes after occurrence. If hypoxemia can be predicted or anticipated before it occurs, then actions can be taken by anesthesiologists to proactively prevent hypoxemia and minimize patient harm.

Machine learning (ML) techniques use statistical methods to infer relationships between patient features and outcomes in large datasets, and have been successfully applied to predict adverse events in health care settings, such as sepsis, or patient deterioration in the intensive care unit (*9–12*). Yet ML techniques to predict adverse events such as hypoxemia in a considerably more critical and complex setting such as the operating room are currently lacking. Moreover, though previous complex ML approaches provide good prediction accuracy, their application in an actual clinical setting is limited because their predictions are difficult to interpret, and hence not actionable. Interpretable methods explain *why* a certain prediction was made for a patient, i.e., specific patient characteristics that led to the prediction. This lack of interpretability has thus far limited the use of powerful methods such as deep learning and ensemble models in medical decision support.

We present an ensemble model based machine learning method, *Prescience*, that predicts the near-term risk of hypoxemia during surgery *and* explains the patient and surgery specific factors that led to that risk (Figure 1). We believe this is an important step forward for machine learning in medicine because while machine learning models have significantly improved the ability to predict a patient’s future condition (*13*), the inability to explain the predictions from accurate, complex models is a serious limitation. Understanding what drives a prediction is important for determining targeted interventions in a clinical setting. For this reason, machine learning methods employed in clinical applications avoid using complex, yet more accurate models and retreat to simpler interpretable (e.g., linear) models at the expense of poorer accuracy. We demonstrate how to retain interpretability, even when complex models such as nonparametric methods or deep learning are used, by developing a method to provide theoretically justified explanations of model predictions building on recent advances in model-agnostic prediction explanation methods (*14–16*). This allows these accurate, but traditionally hard to interpret, models to be used while still providing intuitive explanations of what led to a patient’s predicted risk. Our ability to provide simple explanations of predictions from arbitrarily complex models eliminates the typical accuracy vs. interpretability tradeoff, thus allowing broader applicability of machine learning to medicine.

Prescience was trained to use standard operating room sensors to predict hypoxemic events in the near future and explain why an event is, or is not, likely to occur. It departs from the relatively few previous approaches to this problem in two important ways: First, unlike ElMoaqet et al. who used a linear autoregressive support vector machine on arterial oxygen saturation times series (*9*), and Tarassenko et al. who used Parzen windows to find outliers from five input patient measurement types (*17*), Prescience integrates a comprehensive dataset from a hospital’s Anesthesia Information Management System (AIMS) (see Methods for details). The AIMS data consists of high fidelity *real-time data* – such as time series data from patient monitors and anesthesia machines, bolus and infusion medications, input and output fluid totals, laboratory results, templated and free text descriptions of anesthesia techniques and management, and *static data* – such as American Society for Anesthesiology (ASA) physical status, surgical procedure and diagnoses codes (*18*), as well as patient demographic information such as age, sex, smoking status, height and weight. By continuously integrating a broad set of patient and procedure *features* extracted from the AIMS data, Prescience surpasses human-level accuracy while maintaining consistent performance during every minute of a surgery.

Second, Prescience explains why a prediction was made, regardless of the complexity of the machine learning model used to make the prediction. Significant progress has been made recently integrating predictive machine learning solutions into medical care (*9–12*). However, accurately and intuitively conveying to doctors *why* a prediction was made remains a key challenge. For example, a numeric representation of risk (e.g., 2.4 odds ratio in Figure 1) is useful. However, a more detailed presentation that shows the risk is due to the patient’s BMI, current tidal volume, and pulse rate is more clinically meaningful since some factors may be modifiable and result in clinical changes mitigating that risk.

**Fig 1.**
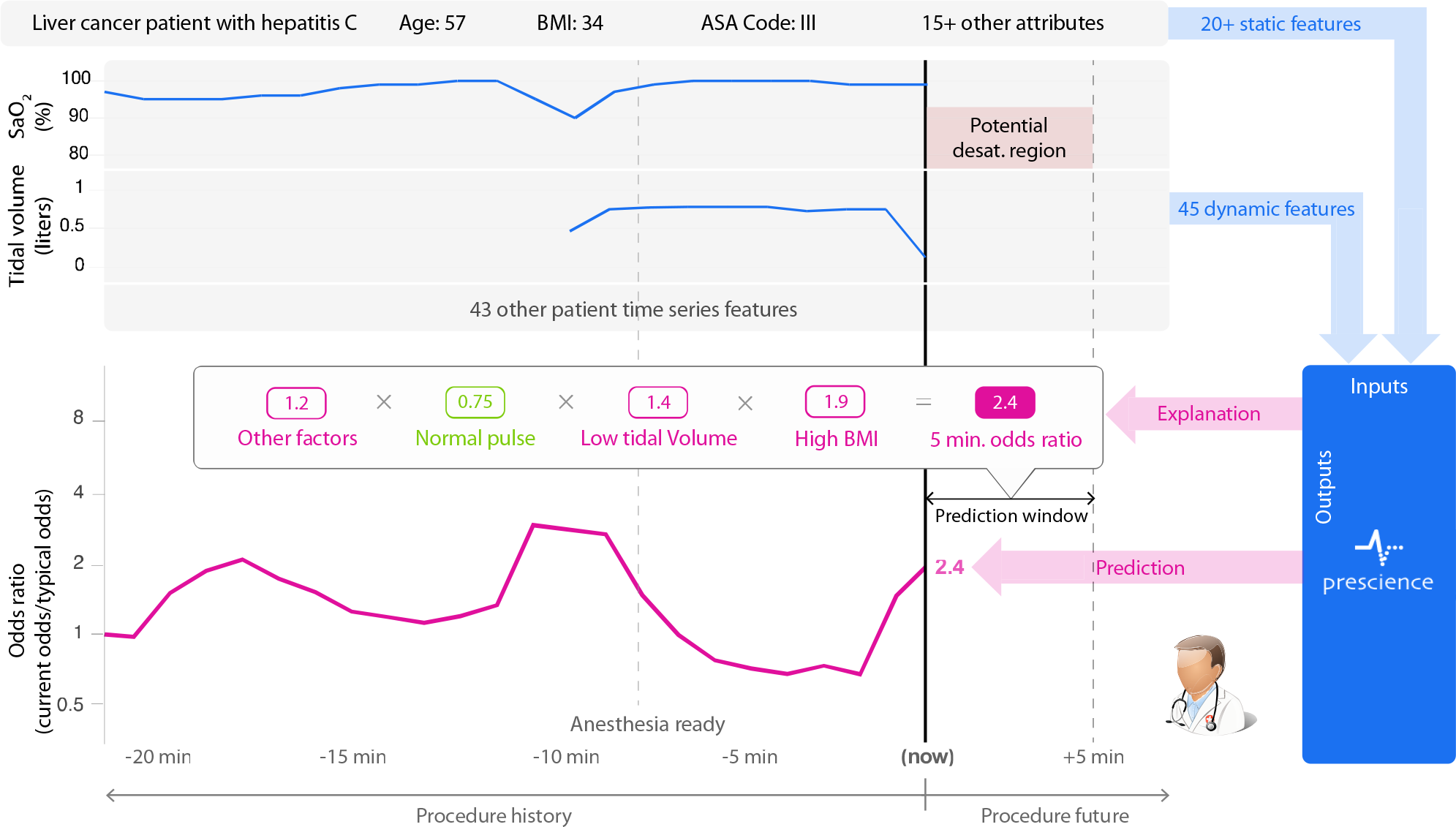
Prescience integrates many data sources into a single risk, which is explained through a succinct visual summary. A wide variety of data sources were used to build a predictive model of hypoxemia events. An explanation (shown above) is then built for each prediction. Purple features have values that increased risk, while green features decreased hypoxemia risk. The combination of impacts of all features is the predicted Prescience risk; in this case the odds are 2.4 times higher than normal.

## Results

To demonstrate the value of Prescience’s explained predictions and gain insight into factors affecting intraoperative hypoxemia, we present the following results: 1) a comparison of Prescience hypoxemia predictions against anesthesiologists’ predictions with and without the aid of Prescience, 2) an example of how Prescience explains hypoxemia risk at a specific time-point during a surgical procedure, 3) a comparative summary of relevant AIMS data features for hypoxemia prediction chosen by Prescience and by anesthesiologists, and 4) a detailed presentation of key risk factors for hypoxemia identified by Prescience.

### Prescience overview – data preparation, model learning and feature importance estimation

Based on World Health Organization recommendations and for the purposes of prediction, we defined hypoxemia as the drop in SpO_2_, i.e., arterial blood oxygen saturation as measured by pulse oximetry, to 92% or lower (see Methods; Supplementary Figure 1). From the AIMS data, we extracted 3,797 *static features* for each patient and an expanded set of 3,905 *real-time* and *static features* for each time point during surgery (see Methods; Supplementary Table 1). We excluded cases (heart transplant, lung transplant, tracheostomy, and coronary artery bypass surgeries) in which SpO_2_ and other hemodynamic parameters can be significantly affected by non-physiological measurements such as during cardiopulmonary bypass. All the experiments were performed after appropriate Institutional Review Board (IRB) approval (see Methods), with clinical data summarized in Table 1.

**Table 1.**
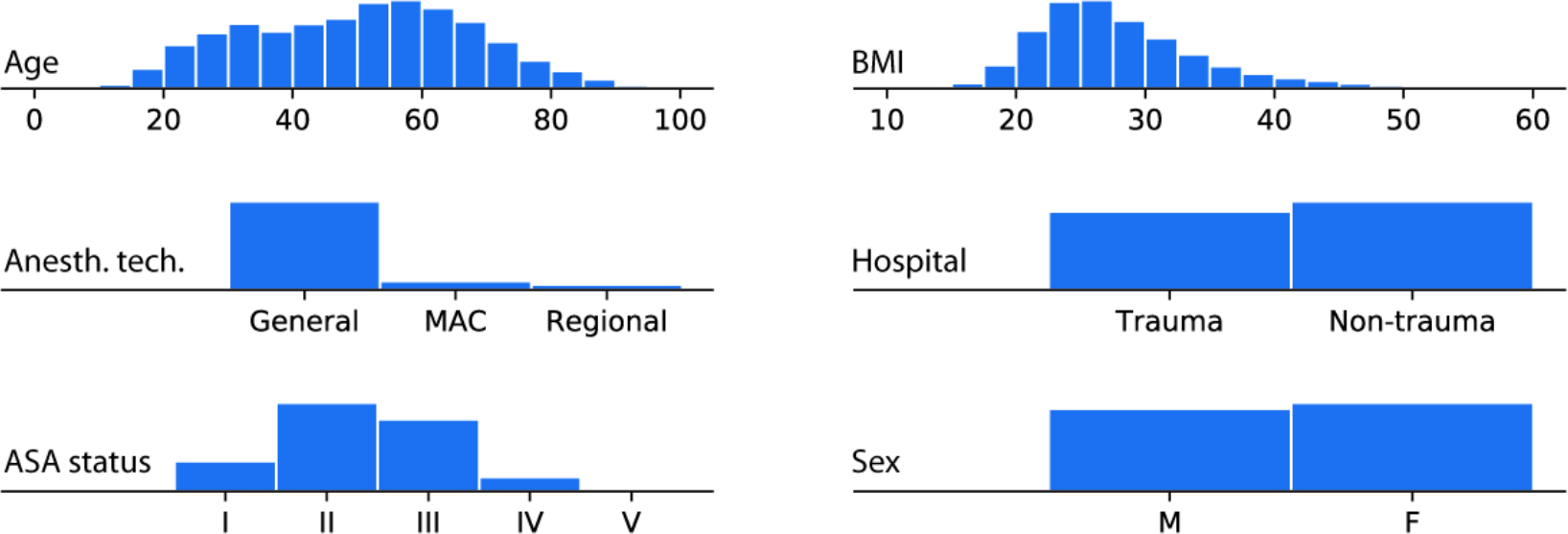
Patient and procedure characteristics. Histograms summarizing basic properties of the anesthesia procedures used for training (ASA stands for American Society of Anesthesiologists, and BMI means Body Mass Index). Prescience was trained and evaluated using data from 53,126 procedures recorded at two hospitals over two years.

We trained a gradient boosting machine model (*19*) to solve the following two types of prediction problems: A) *initial prediction*: predicting at the start of a procedure the risk of hypoxemia anytime during a procedure based on the static features, and B) *real-time prediction*: predicting in the next 5 minutes at various points of the operative period based on real-time and static features collected up to that time point. For task A) we used 42,420 procedures as *training samples* to train the gradient boosting machine, 5,649 procedures as *validation samples* to choose the tuning parameters for the gradient boosting machine (and other prediction models for comparison), and 5,057 as *test samples* for comparing across different prediction models (Supplementary Figure 4). For task B), we used 8,087,476 time points as training samples, 1,053,629 as validation samples, and 963,674 as test samples, where all time points from the same procedure were included in the same sample set (Supplementary Figure 3). To ensure that there was no bias towards the final test set, the test data was initially compressed and left compressed until method development was completed.

As shown in Supplementary Figures 3 and 4, the gradient boosting machine outperforms alternative prediction models previously used for similar problems.

For tasks A) and B), we use 198 and 523 test samples respectively for evaluating anesthesiologists’ performance for initial and real-time prediction tasks, respectively (see below). Prescience outputs the risk prediction and its explanations (Figure 1; Figure 3A) which show a set of features that increased (purple-colored) and decreased (green-colored) the risk.

We developed an efficient, theoretically justified machine learning technique to estimate the importance of each feature on a prediction made for a single patient, which drives real-time explanations (Figure 3) for the Prescience model. We verified the quality of the explanations given to the anesthesiologists (in the experiments described below) by comparing the explanations with the change in model output when a feature is perturbed (Supplementary Figure 5). We also developed effective visualizations of these explanations that encodes them in a compact visual form for anesthesiologists (Figure 1; Supplementary Figures 6-8), and a more detailed visualization which highlights the relevant contributing features (Figure 3) (see Methods for details).

### Prescience improves anesthesiologist’s ability to predict hypoxemia

To test the potential of Prescience to aid hypoxemia prediction we replayed prerecorded surgery data from test sample procedures in a web-based visualization to five practicing anesthesiologists (Supplementary Figures 6-8). Each anesthesiologist was given two types of prediction tasks: A) *initial prediction* (198 tasks), and B) *real-time prediction* (523 tasks). For each prediction task, anesthesiologists were asked to provide a *relative risk* of hypoxemia as compared to a normal acceptable risk, for example, 0.01 for 1/100^th^ the normal risk or 3.4 for 3.4 times the normal risk. These relative risks were then used to calculate standard receiver operating curves averaged over five anesthesiologists as shown in Figure 2 which plots the true positive rate (i.e., % of desaturations correctly predicted) in the y-axis against the false positive rate (i.e., % of non-desaturations incorrectly predicted) in the x-axis.

Figure 2A-B shows that for both types of prediction tasks, predictions made by Prescience (purple) are considerably more accurate than anesthesiologists’ predictions (green). The prediction accuracy of anesthesiologists (green) markedly improved when the anesthesiologists were given Prescience’s risk prediction and its explanations in addition to the original procedure data (blue) (Supplementary Figures 6-8). A clear separation between the performance of anesthesiologists with (blue) and without (green) the aid of Prescience is observed for both initial prediction (Figure 2A, P-value < 0.0001) and real-time prediction (Figure 2B, P-value < 0.0001). This suggests that Prescience can enhance anesthesiologists’ assessment of future risk and their ability to proactively anticipate hypoxemia events. Interestingly, the prediction performance of anesthesiologists with Prescience explanations (blue) was slightly lower than direct predictions from Prescience (purple). This means that when the anesthesiologists adjust their risk estimate for a patient away from what Prescience originally predicted they are more likely to be wrong than right.

**Fig 2.**
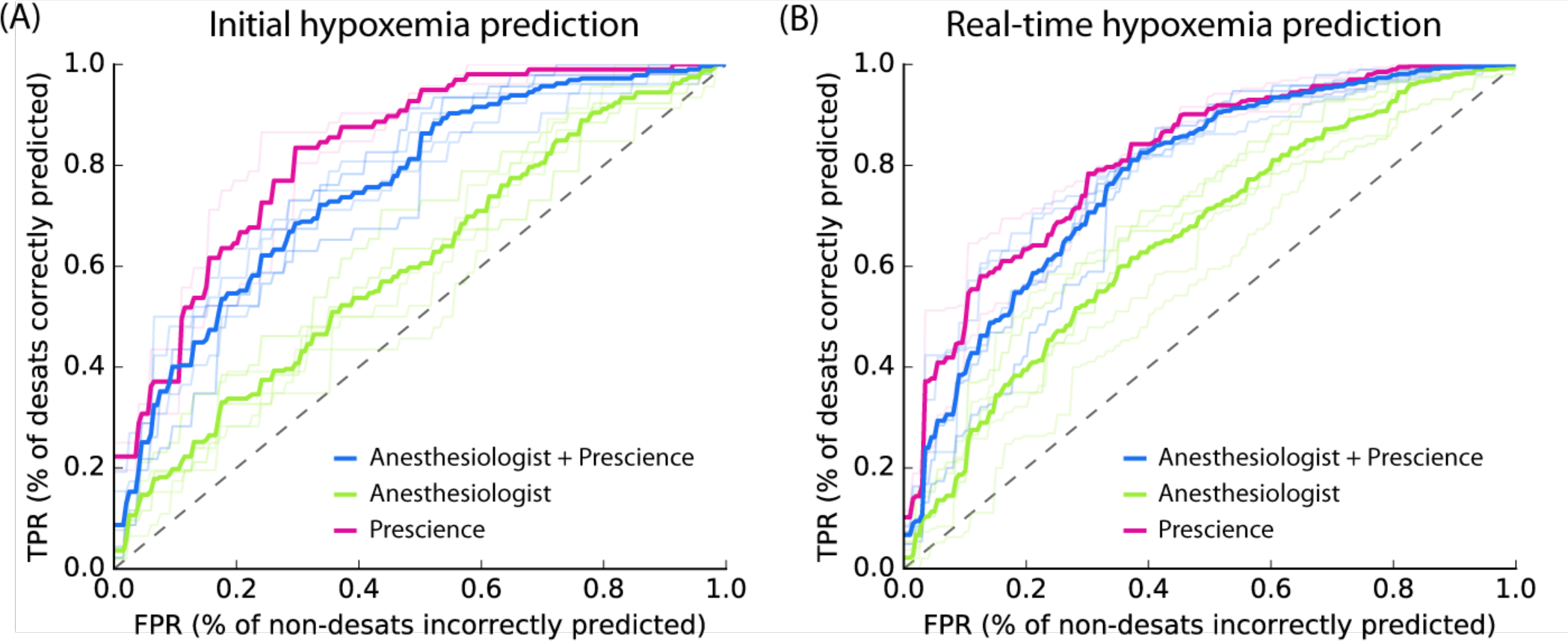
Pooled comparison of five anesthesiologists’ prediction performance with and without assistance by Prescience. Receiver Operating Characteristic (ROC) plots comparing five anesthesiologists’ predictions from recorded data with and without Prescience assistance. Light colored lines represent individual anesthesiologist’s performances; dark lines represent their average performance. (**A**) For initial risk prediction, anesthesiologists (green, AUC = 0.60) performed significantly better with Prescience assistance (blue, AUC = 0.76; P-value < 0.0001) than without Prescience assistance, and Prescience performed better in a direct comparison with anesthesiologists (purple AUC = 0.83); P-value = 0.0003). (**B**) For intraoperative real-time (next 5 minute) risk prediction anesthesiologists (green, AUC = 0.66) again performed better with Prescience assistance (blue, AUC = 0.78; P-value < 0.0001), and Prescience alone outperformed anesthesiologists predictions (purple, AUC = 0.81); P-value = 0.0068). Note that The False Positive Rate (FPR) (x-axis) measures how many points without upcoming hypoxemia were incorrectly predicted to have upcoming hypoxemia. The True Positive Rate (TPR) (y-axis) measures what percentage of hypoxemic events was correctly predicted. P-values were computed using bootstrap resampling over the tested time points.

To avoid the scenario in which an anesthesiologist is tested on the same prediction task twice – one with and the other without Prescience, we created replicate test sets by dividing the prediction tasks into two groups of similar size: (100, 98) tasks for initial prediction and (260, 263) tasks for real-time prediction. Each of the five recruited anesthesiologists was assigned to receive Prescience’s assistance in one of these two replicate test sets (Methods). The procedures shown to anesthesiologists were chosen such that ~50% showed at least one incident of hypoxemia (for preoperative prediction), and time points were chosen such that ~33% had hypoxemia in the next 5 minutes (for intraoperative prediction).

If we extrapolate the real-time results to the 30 million annual surgeries in the US under the assumption that doctors anticipate 15% of hypoxemic events while SpO_2_ is still ≥ 95, then with Prescience assistance they may be able to anticipate 30% of these events, or approximately 2.4 million additional episodes of hypoxemia annually (defined here as SpO_2_ ≤ 92). Since 20% percent of the Prescience risk prediction is based on drugs and settings under the control of the anesthesiologist (Supplementary Table 5; Supp_RealtimeFeatureTable_5.csv), a large portion of these predicted events may be preventable.

### Explained risks reveal both procedure and time specific effects

An explanation from Prescience represents the effects of interpretable groups of patient features (see Figure 1 and Figure 3A). These effects explain why the model predicted a specific risk, and thus allow an anesthesiologist to plan appropriate interventions. In Figure 1 only the most significant features contributing to hypoxemia risk are shown for quick reference, however in Figure 3 the relative contributions of all patient and case features (i.e. attributes) towards hypoxemia risk can be seen at every sample time point during a procedure (Figure 3B). Without a meaningful explanation, the sudden increase in risk shown at the time point marked ‘Now’ might be hard to interpret; however, by representing the predicted risk as a cumulative effect of contributing patient and procedure features, the reason for the increase becomes clear (Figure 3A).

The increase in the risk of hypoxemia in the next 5 minutes shown in Figure 3 is driven by a set of features capturing both static attributes, such as patient height and weight, and dynamic parametric values, such as tidal volume (i.e., volume of gas exhaled per breath) and administration of drugs. The risk explanation bar in Figure 3A has purple-colored features that push the risk higher (to the right) and green-colored features that push the risk lower (to the left). Each group of features is sorted by the magnitude of their impact, and the largest impact features are labeled. Through this representation we can see that many of the 3,905 real-time features have only a small impact, and the risk for this time point is predominantly driven by a few features. Figure 3B shows the trend in the Prescience risk predictions over the course of the procedure. The plot in Figure 3B is equivalent to rotating the feature explanation in Figure 3A by 90 degrees and then stacking the explanations for each time point horizontally. We can see from the risk trend in Figure 3B that the large increase in risk at the current time was driven by ‘Tidal volume’, meaning a drop in the patient’s tidal volume. The future SpO_2_ (blood oxygen concentration) measurements confirm that the patient did indeed progress to hypoxemia (i.e., SpO_2_ ≤ 92). Not only does Prescience alert anesthesiologists when a patient’s risk for hypoxemia is high, but also provides information on the factors and their relative contributions driving the risk. This informed risk prediction enables anesthesiologists to plan an appropriate course of action to avoid hypoxemia.

**Fig 3.**
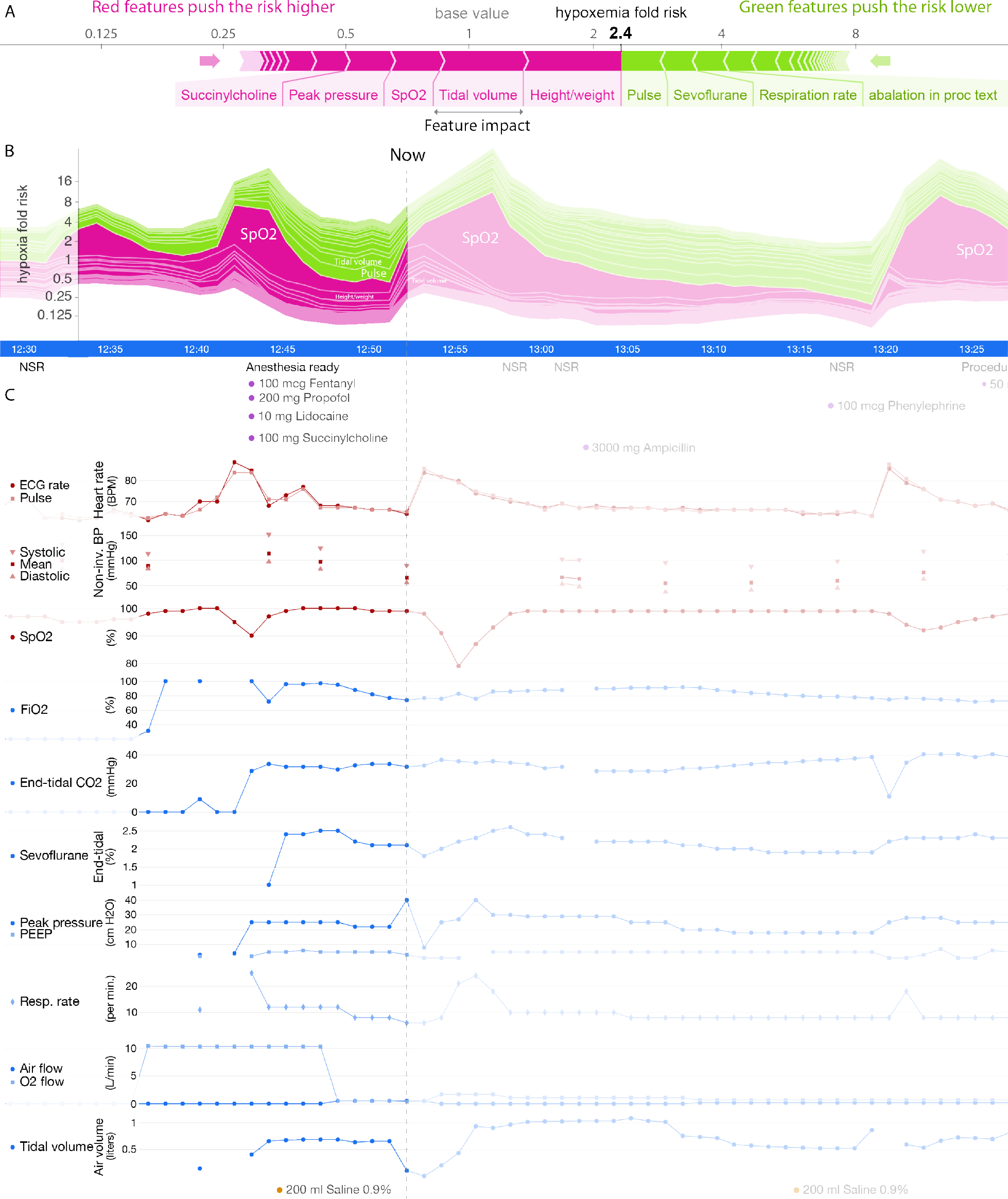
Sample real-time prediction during a procedure. One hour of data is shown from a procedure. (**A**) Explained risk of hypoxemia in the next five minutes. (**B**) Plot of the explained risks evolving over time. This plot is equivalent to rotating (A) 90 degrees and stacking the risk explanations for every time point horizontally. (**C**) A subset of the patient data for this procedure, plotted both before and after the current time point.

### Averaged feature importance estimates broadly align with a survey of prior expectations

To gain an understanding of the general impact of features across all procedures we computed the average importance of each feature in the Prescience model. In contrast to the explanations shown in Figure 1 and Figure 3A which are specific to a single prediction at a particular time point, these average feature importance estimates are over many procedures and time points (*20*). These averaged feature importance estimates are shown for both initial prediction (Figure 4A) and real-time prediction (Figure 4B).

To estimate which clinical features anesthesiologists use to estimate hypoxemia risk, we first performed a survey before using Prescience, which asked four anesthesiologists to list the most important factors they consider when assessing the risk of hypoxemia, both before (for initial prediction) and during a procedure (for real-time prediction). Their responses were then aggregated into a single ranked list of features (Supplementary Tables 2 and 3). Figure 4 shows the rankings chosen by anesthesiologists next to the feature importance estimates derived by Prescience for (A) initial and (B) real-time predictions. The ranking of features by anesthesiologists appears to correspond well with the ranking by Prescience.

As another way to measure which features anesthesiologists’ think contribute to hypoxemia, we learned from anesthesiologists’ behavior by training a separate gradient boosting machine model based on their predictions. This allows a direct comparison between the anesthesiologists and Prescience on the same set of features. We fit this model to all the anesthesiologist relative risk predictions using 10-fold cross validation. We then computed the feature importance estimates for this model that was trained to mimic the behavior of anesthesiologists. Given the relatively smaller set of training examples used to train the model (198 initial predictions, and 523 real-time predictions), we used bootstrapping to estimate the variability of the feature importance estimates (Figure 4 right).

In general, there is reasonable agreement between the Prescience feature importance estimates and those identified by the anesthesiologists. However, there are important differences that may stem from the comprehensive nature of the Prescience analysis, while anesthesiologists necessarily focus on what they consider the most likely causes for hypoxemia concern. One striking difference is the reduced role of current SpO_2_ levels in anesthesiologists’ predictions. While anesthesiologists are clearly influenced by the recent patterns of patient SpO_2_ levels, Prescience strongly depends on these patterns, while anesthesiologists appear to be equally influenced by other factors, such as end tidal CO_2_ and peak ventilation pressure.

Our study used data from two hospitals and initial hypoxemia predictions were driven by a bias between the two hospitals. This is perhaps unsurprising since one hospital is a level 1 trauma center and a significant proportion of its surgical cases involve trauma patients who are more susceptible to hypoxemia. However, it is interesting to note that the importance of hospital as a risk factor became insignificant for the intraoperative real-time predictions, presumably because the risk differences in each hospital were captured by the real-time features.

Among the static features, BMI (body mass index) and age were significant risk factors. These features are well understood in the medical literature as risk factors that can increase the chances of hypoxemia (*21, 22*). The American Society of Anesthesiology (ASA) physical status feature represents the severity of a patient’s medical condition and a higher ASA number indicates a higher comorbidity. Prescience determined that higher ASA physical status values predisposes a patient to higher hypoxemia risk. While this finding may be clinically intuitive, anesthesiologists can now use this information in their preoperative evaluation as a pre-specified risk factor for intraoperative hypoxemia. Eye procedures were informative to the model and carried a reduced risk of hypoxemia, while surgeries for fractures had a slightly higher risk. These patterns may reflect the composite risk of hypoxemia to patients undergoing these particular procedures. In the case of eye surgeries, the risk was lower even though many are elderly and have accompanying co-morbid conditions. The low risk of eye procedures is in contrast to fractures which carry an increased risk for hypoxemia. Together these findings provide new data on the relative “risk” of these procedures which has implications for anesthesia staffing, need for equipment, and preparation for the ability to rescue patients from hypoxemia. Eye procedures and surgeries for fractures are two examples of text based features extracted from diagnosis and preoperative procedure notes. They demonstrate that unstructured text notes can be combined with structured patient data to improve patient risk prediction. Although many of the risk factors identified by Prescience reconfirmed those expected by the anesthesiologists, it is informative that Prescience independently identified these features with no prior knowledge.

Among real-time (intraoperative) features SpO_2_ (arterial oxygen saturation) is, as expected, the strongest predictor of future potential drops in SpO_2_. End tidal CO_2_ (amount of carbon dioxide exhaled by the patient) was also a significant intraoperative feature identified by Prescience as predictive of hypoxemia. Lower values may indicate inadequate ventilation or airway obstruction which can turn increase the risk of hypoxemia. Prescience also determined that hypotension (systolic blood pressure below 80) increases the risk of hypoxemia. On the other hand, increased FiO_2_ (inspired O_2_ concentration) and adequate positive pressure ventilation can reduce the risk of hypoxemia, as expected by the anesthesiologists.

**Fig 4.**
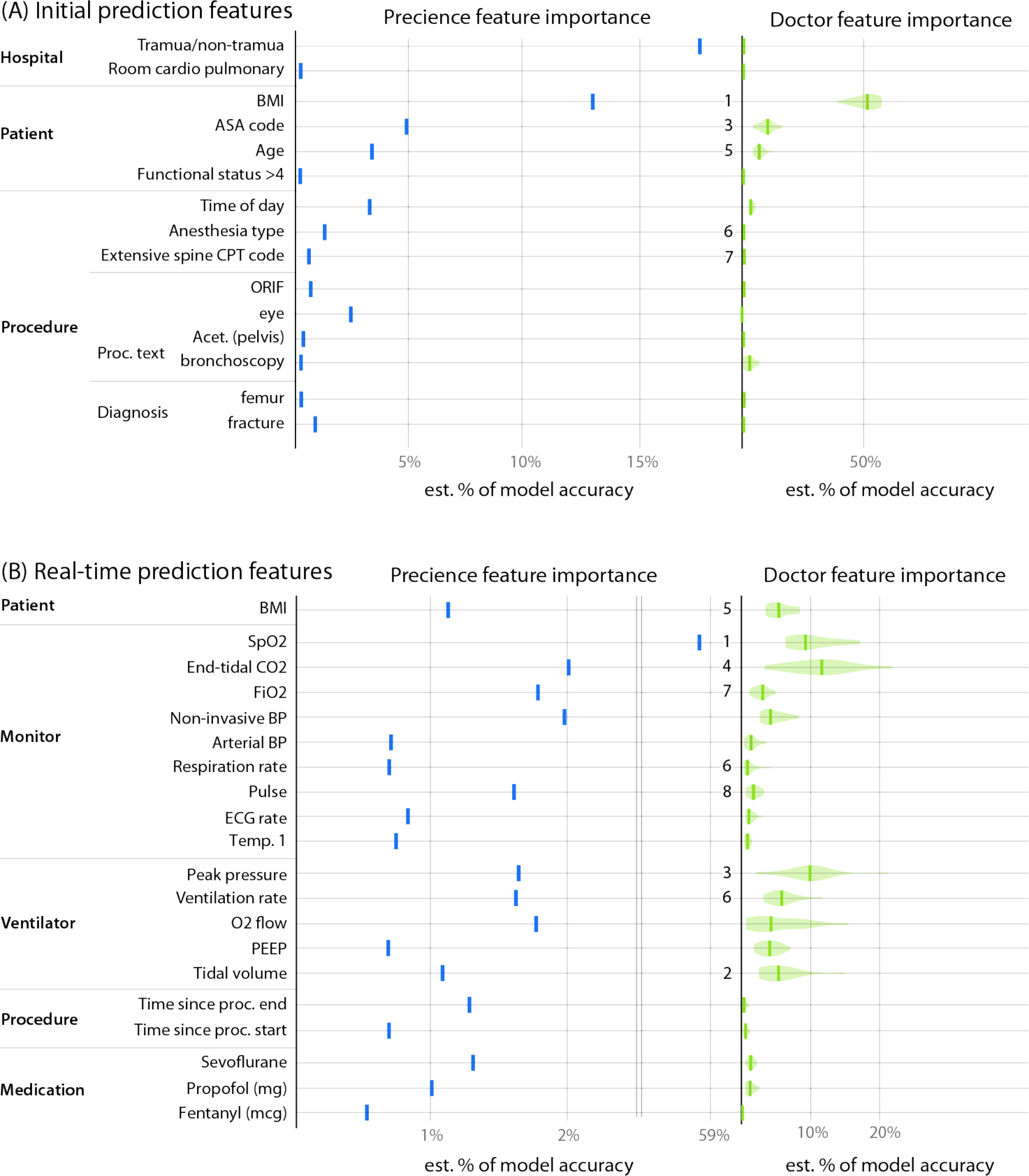
Comparison of averaged feature importance estimates between Prescience and anesthesiologists for both initial and real-time prediction. Importance estimates assigned by the Prescience model and anesthesiologists to the top features in both (**A**) initial and (**B**) real-time prediction. The importance of features is measured as the estimated percent of the model’s prediction accuracy that is due to that feature. The numbers presented to the left of the imputed anesthesiologist importance estimates are feature rankings from a consensus of anesthesiologist responses about which features they believed would be important. The quantitative anesthesiologist feature importance estimates were estimated using 20 bootstrapped models trained to mimic the anesthesiologist’s predictions when unassisted by Prescience.^1^

### Prescience’s estimated importance of individual features on hypoxemia risk highlight important clinical relationships

Three important features each for beginning of surgery (initial) and during surgery (real-time) predictions were chosen to illustrate how the Prescience model modifies hypoxemia risk based on changes to feature characteristics (Figure 4). While many such relationships are present for the various features, Figure 5 shows a representative selection demonstrating informative risk relationships that are captured in the Prescience model.

Among static features we find that patient BMI has a clear effect on the risk of hypoxemia. When the BMI is over 26, the risk of hypoxemia increases linearly until it has more than doubled when BMI is over 50. Though a qualitative association between hypoxemia and body weight is well established in the field of anesthesia (*21, 22*), Prescience quantifies this relative risk.

Prescience shows that patients with higher ASA physical status codes have higher risk of intraoperative hypoxemia. This is not surprising since higher ASA codes represent increased severity of a patent’s physical condition such as preexisting pulmonary and cardiac conditions that can predispose a patient to develop hypoxemia. Prescience data support clinical observations that the risk of hypoxemia more than doubles when the ASA code increases from I to V. Advancing age also predicted intraoperative hypoxemia, likely representing the presence of comorbidities (*21*). These data show that BMI > 30, which meets the clinical definition of obesity (*23*), is associated with intraoperative hypoxemia, suggesting impaired pulmonary mechanics. While we agree that these findings confirm clinical observations and suspicions of the relationship between these patient factors and adverse anesthesiology outcomes, Prescience quantifies this association and the risks, giving a more clinically useful interpretation to anesthesiologists.

For real-time prediction, measurements from each time series are represented by a set of multiple features. For simplicity, we focus here only on the effect of the shortest time lag exponentially weighted moving average, which essentially represents the most recent reported value in the time series (see Methods for details).

Tidal volume represents the amount of gas exhaled per breath when the patient is either breathing spontaneously or mechanically ventilated during general anesthesia. As the tidal volume drops below 0.6 liters (keeping all other features the same), Prescience risk for hypoxemia increases. This increase could be due to hypoventilation, in which case anesthesiologists take preventative steps to avoid inadequate ventilation.

End tidal CO_2_ represents the amount of carbon dioxide exhaled gas. Figure 5 shows the relationship between end tidal CO_2_ and risk of hypoxemia under general anesthesia. End tidal CO_2_ below 35 mmHg is associated with an increasing risk of intraoperative hypoxemia. While we cannot definitively attribute hypocapnia with intraoperative hypoxemia, these associations may represent underlying patient conditions such as chronic obstructive pulmonary disease that affect both physiological conditions. Alternately, the low end tidal CO_2_ and may result from either intentional or unwanted hyperventilation during anesthesia care.

Examining FiO_2_ is important because anesthesiologists can control the amount of oxygen delivered to patients. Current practice is to not provide all patients with 100% FiO_2_ since not all patients need it, prolonged ventilation with 100% FiO_2_ is associated with pulmonary atelectasis, and because oxygen if delivered when not needed is costly and wasteful. These data show that FiO_2_ below 40% is independently associated with intraoperative hypoxemia irrespective of other features. These findings provide important information regarding safe practice of FiO_2_ in patients during general anesthesia. It is possible that the routine practice of maintaining FiO_2_ 30% or close to room air may be harmful to patients and not desirable. While these effects are adjusted for all other available features, it is important to note that as with any observational study some residual confounding with patient risk may still exist. This could explain the increase in hypoxemia risk we observed for high O_2_ levels.

These representative features illustrate the power of our machine learning-based prediction method, Prescience, to not only provide explained risk predictions, but also quantitative insights into the exact change in risk induced by certain patient or procedure characteristics.

**Fig 5.**
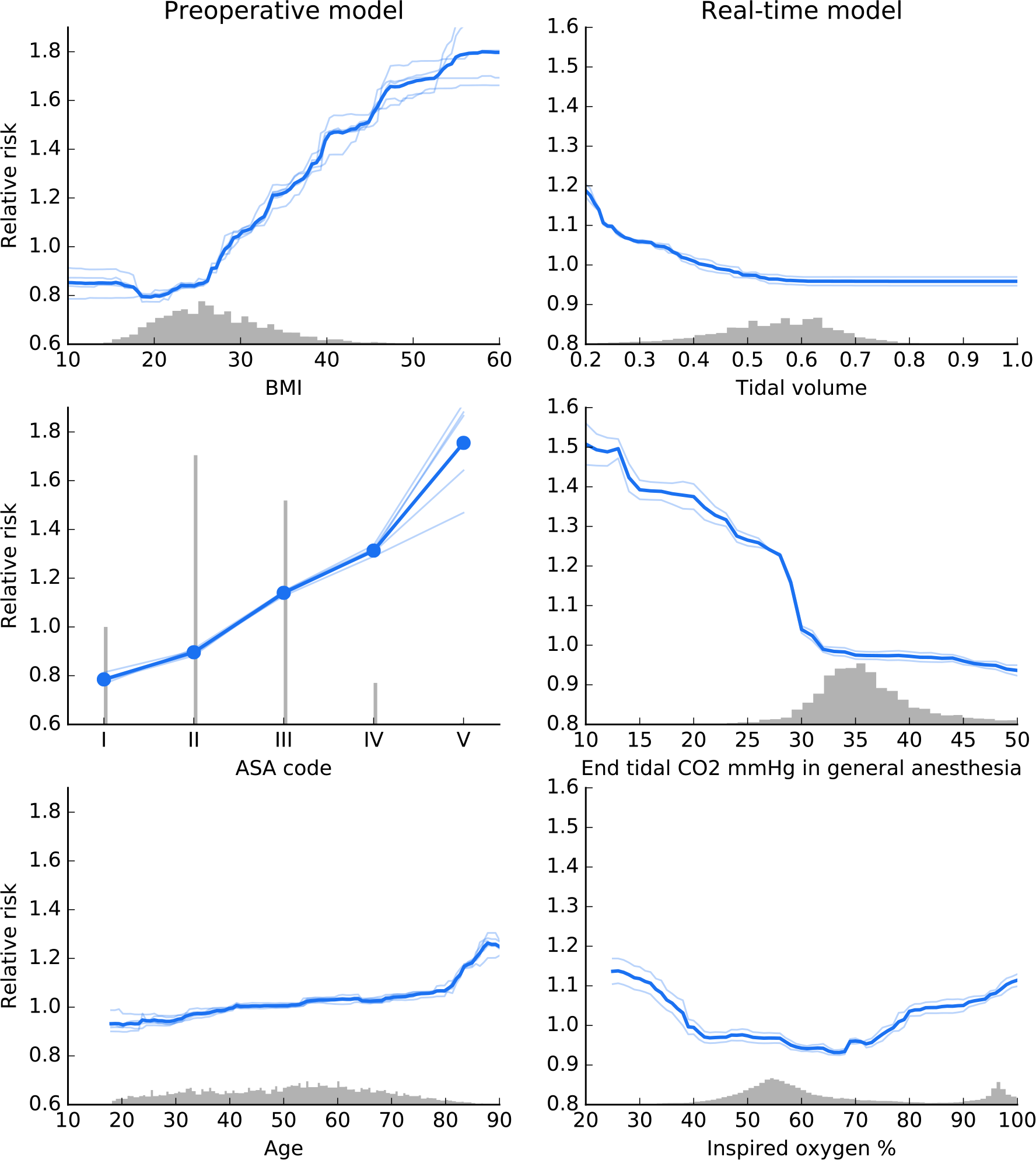
Effect of varying individual feature values for both initial features (left) and real-time features (right). These partial dependence plots show the change in hypoxemia risk for all values of a given feature. The gray histograms on each plot show the distribution of values for that feature in the validation dataset. Light colored lines represent model variability from bootstrap resampling of the training data.

## Discussion

To the best of our knowledge, Prescience is the first method designed to comprehensively integrate high fidelity operating room data to predict intraoperative hypoxemia events before they occur. Based on a comparison against practicing anesthesiologists and existing computational methods applied to other clinical problems, Prescience achieves superior performance when predicting hypoxemia risk from electronically recorded data.

To our knowledge, Prescience is also the first method to combine high accuracy complex models with interpretable explanations. This combination of accuracy and interpretability allows physicians to receive the best possible predictions while also gaining insight into why those predictions were made. To test how Prescience predictions with explanations would impact an anesthesiologist’s ability to estimate hypoxemia risk we compared anesthesiologist predictions with and without Prescience assistance. We observed a clear increase in prediction accuracy when doctors were assisted by Prescience, demonstrating that anesthesiologists may make more accurate hypoxemia risk assessments in the operating room if they had access to Prescience.

It should be clarified that our exercise at developing machine learning methods to predict intraoperative hypoxemia, though promising, should still be considered an initial attempt. In this first attempt, we did not categorize procedures to assess hypoxemia predictions in specific types of procedures. For this reason, clinical interpretation of the results had to be somewhat generic. For enhanced interpretation of risks, future attempts need to focus on specific categories of cases and phases of anesthesia. Another future enhancement would be integrating additional preoperative data such as patient’s detailed medical history into the prediction models. Higher fidelity intraoperative data such as patient monitor waveform data could enrich machine learning thus potentially leading to more accurate predictions. Prospective trials of Prescience during live procedures are also needed to verify the improvements in anesthesiologist’s performance we retrospectively observed in prerecorded procedures.

This paper focuses on hypoxemia risk during intraoperative anesthesia care. However, the importance of coupling accurate predictions from complex models with interpretable explanations of why a prediction was made, has broad applicability throughout medicine. Extending the approach taken by Prescience and providing these model explanation tools to the community is a clear next step. Because Prescience effectively decouples the interpretable explanation from the prediction model, we are also free to continue to refine the core prediction model without changing the user experience for anesthesiologists.

The global risk profiles learned by Prescience (Figures 4-5) are clinically relevant for a number of reasons. First, they show that in the health system examined, trauma hospital patients may be more critically ill as they have more intraoperative hypoxemia. In current times when harmonization of care and standardization are considered to reduce unwanted clinical variation, these data suggest that resources may need to be differentially deployed to address differential rates of adverse events. Second, anesthesiologists can now quantify risks of intraoperative hypoxemia adjusted for other factors to the very elderly, those who are overweight, and those with more comorbid conditions. The exact relationships described in Figure 5 clearly show the patterns and threshold points for the risk. Whereas low tidal volume is often suggested for patients with acute lung injury (*24*), these data suggest that overall, low lung tidal volumes are, in-fact, associated with intraoperative hypoxemia. The relationship between low end-tidal CO_2_ levels and intraoperative hypoxemia may reflect underlying critical illness. Despite our inability to fully exclude residual confounding, these data shed new light on physiological relationships as well as provide a mechanism to facilitate provision of anesthesia care that can mitigate intraoperative hypoxemia.

The field of medicine is full of many exciting data science challenges that have the potential to fundamentally impact the way medicine is practiced. More and more data driven predictions of patient outcomes are being proposed and used. However, black-box prediction models which provide simply predictions, without explanation, are difficult for physicians to trust and provide little insight about how they should respond. The interpretable explanations used by Prescience represent a powerful technique that can transform any current prediction method from one that provides *what* the prediction is, into one that also explains *why*.

## Materials and Methods

### IRB statement

The electronic data for this study was retrieved from institutional electronic medical record and data warehouse systems after receiving approval from the Institutional Review Board (University of Washington Human Subjects Division, Approval #46889). Protected health information was excluded from the data set that was used for machine learning methods.

### Data sources

Our hospital system has installed an Anesthesia Information Management System (AIMS) (Merge AIM, Merge Inc, Hartland, WI) that automatically captures minute by minute hemodynamic and ventilation parameters from the patient monitor and the anesthesia machine. The system also integrates with other hospital electronic medical record (EMR) systems to automatically acquire laboratory and patient registration information. The automatic capture of data is supplemented by manual documentation of medications and anesthesia interventions to complete the anesthesia record during a surgical episode. For the current project, we extracted the high-fidelity anesthesia data from the AIMS database for the period May 2012 through June 2014. Additionally, for each patient, medical history data were extracted from our EMR data warehouse (Caradigm, Bellevue, WA). The high-fidelity anesthesia record data and the corresponding medical history data from the hospital EMR formed the underlying data for machine learning. The various data elements used for machine learning are outlined in Supplementary Table 1.

**Supplementary Table 1.**
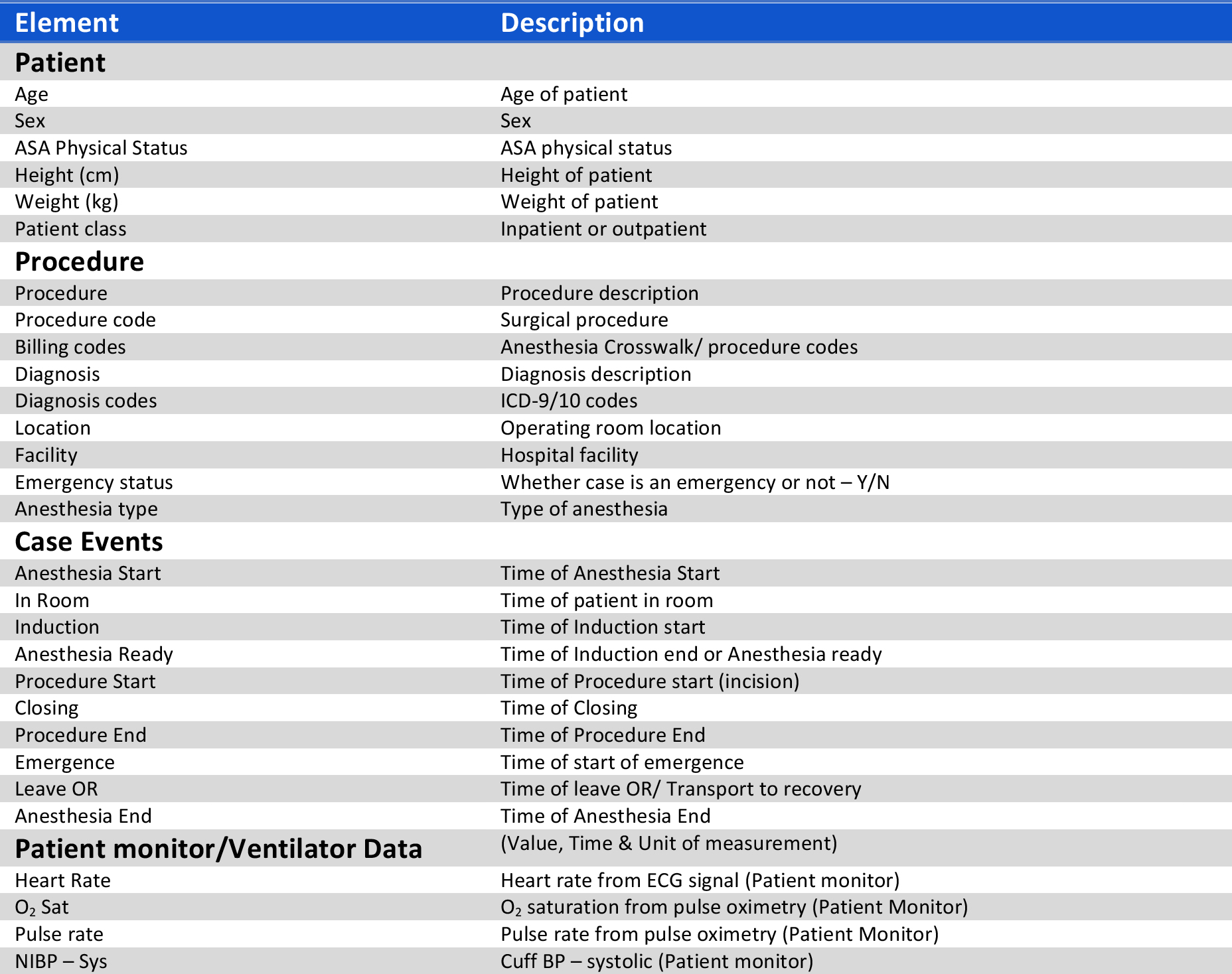
Raw data sources used for machine learning. A wide variety of data sources were used that included text values, time series, discrete values, and quantitative values.

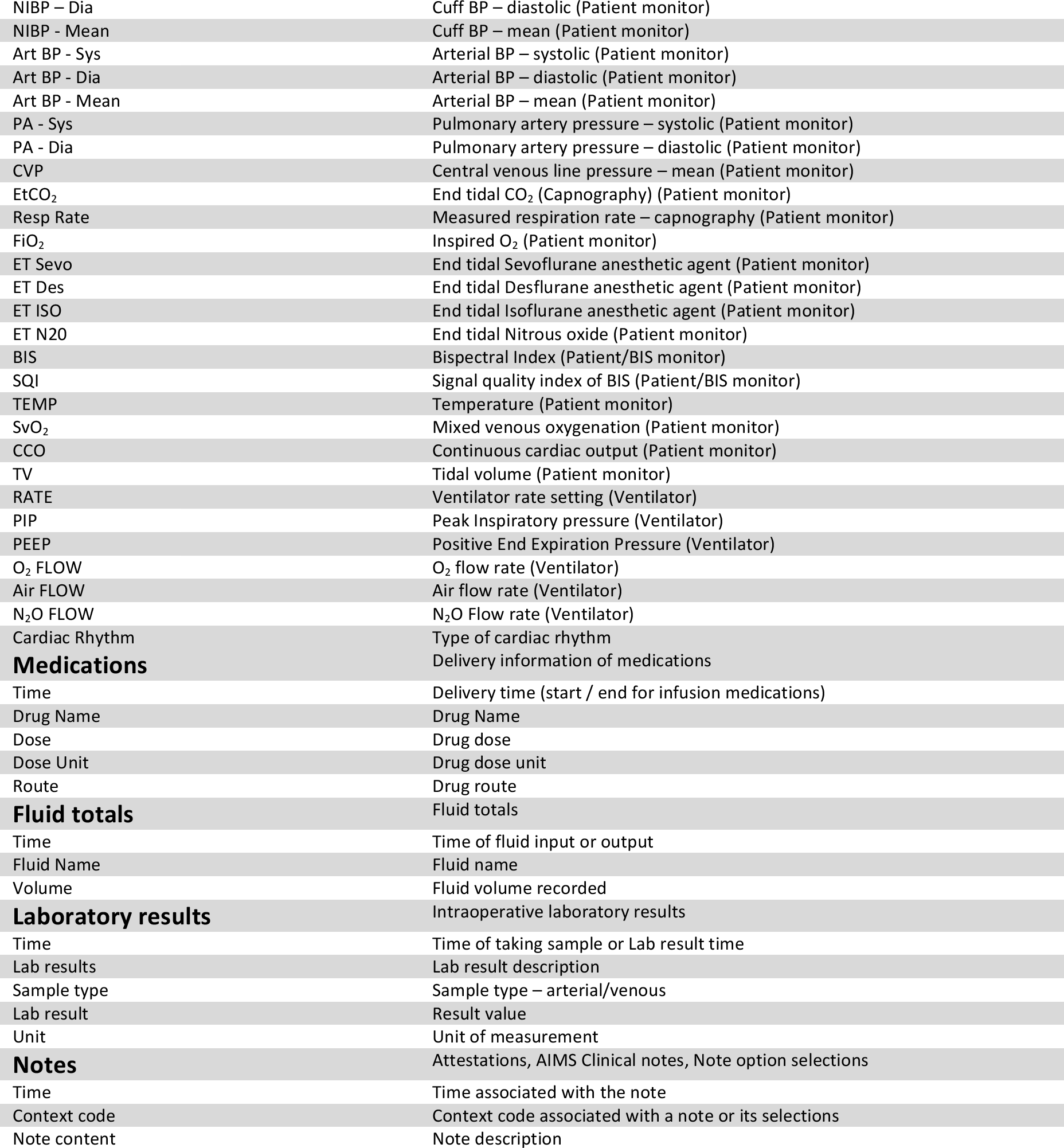

### SpO_2_ desaturation labels

We considered SpO_2_ ≤ 92% as hypoxemia, which falls between the World Health Organization’s recommended intervention level (< 94%) and emergency level (< 90%) (*25*). Predictions of hypoxemia were made for a window 5 minutes into the future. If the SpO_2_ was ≤ 92% at any point during those 5 minutes then it was considered a positive label, otherwise it was negative. The machine learning algorithm was trained using these *training labels* on all time points where SpO_2_ was not already ≤ 92% at that time point.

When evaluating the machine learning algorithm’s performance by comparing with anesthesiologists we chose to use a much more stringent definition of hypoxemia. This more stringent definition excludes some of the time points, and leads to a smaller set of *testing labels. Testing labels* were positive only if SpO_2_ was ≥ 95% for the past 10 minutes and then fell below 92% in the next five minutes (Supplementary Figure 1; left). *Testing labels* were negative only if SpO_2_ remained ≥ 95% for the past ten minutes and the next ten minutes (Supplementary Figure 1; right). All the other cases do not have testing labels. This more restrictive labeling scheme ensures that positive testing labels are clear drops in SpO_2_ levels, while negative testing labels are clearly not drops in SpO_2_ (Supplementary Figure 1).

**Supplementary Fig 1.**
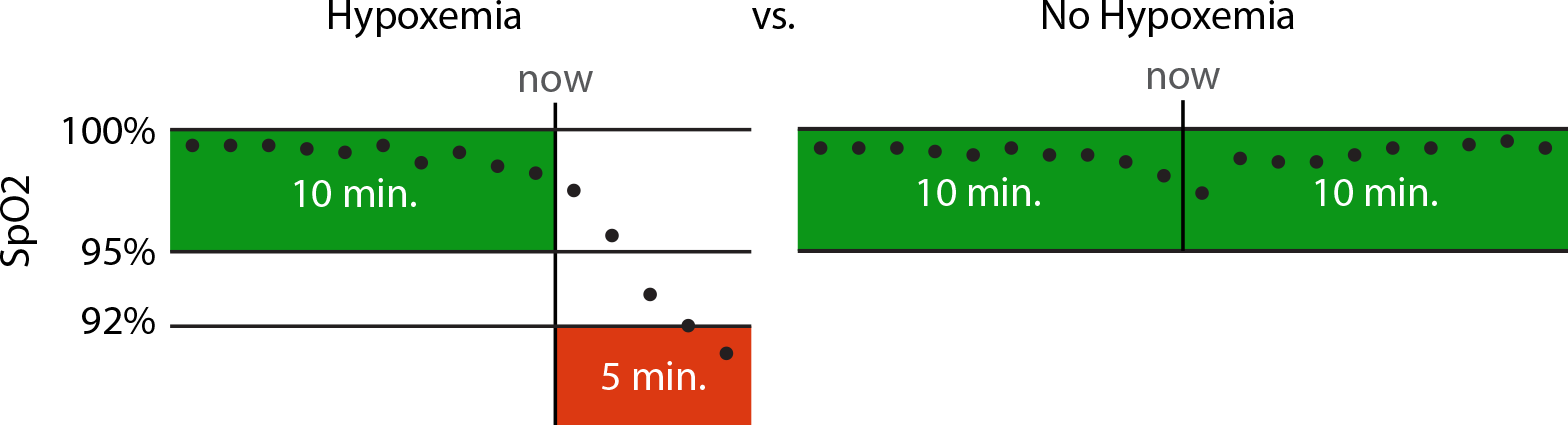
Criteria for defining testing labels. When comparing the performance with anesthesiologists, only time points that clearly either desaturate or not are used. Hypoxemia involves dropping from ≥ 95% to ≤ 92% in the next 5 minutes. Not desaturating means remaining ≥ 95% for both the past 10 minutes and the next 10 minutes.

An important point to consider when building labels for health outcome prediction is that anesthesiologist interventions can affect outcomes. It was recently noted by Dyagilev et al. that models can learn when an anesthesiologist is likely to intervene and hence lower the risk of an otherwise high-risk patient (*26*). This means that patients with low risk (from the model) may still need treatment. To address this, they proposed removing examples from the training set where anesthesiologists have intervened. This allows one to learn a model which predicts patient outcome without intervention. In our case, it is not possible to fully identify when or how an anesthesiologist is intervening (and if that intervention prevented hypoxemia), so we sought to address this issue in two ways:

1. It must be recognized that the model predicts hypoxemia when following standard procedures, *not* the occurrence of hypoxemia if the anesthesiologist takes no action to influence hypoxemia. This is a natural assumption in the operating room where interventions that may affect SpO_2_ levels are performed frequently.
2. By focusing on clear explanations of why a certain risk was predicted we enable anesthesiologists to identify when the algorithm may be basing its risk on their actions vs. when the risk is based on other factors.

### Extracting time series features

To make a prediction at an arbitrary point in time, a consistent set of features should be computed that capture the information present in all previous time points. All the data provided about a procedure is associated with a specific date and time. Text data has the time it was provided, minute-by-minute data from the patient monitor has the time at which each measurement was taken, and single point measurements have the times they were recorded.

We summarized these unevenly sampled time registered data into a fixed length feature vector at any point in time using several complementary methods:

- Patient data, procedure information, and pre-operative notes are represented by a “last value” feature, which is zero before any data is recorded and the data’s value afterwards.
- Time series data is captured using exponentially decaying weighted average and variance estimates using multiple decay rates. These decay rates specify how much impact each past time point has on the computed mean or variance for the time series. We used 6 second, 1 minute, and 5-minute half-life times to capture both high and low frequency components of the signal in each time series (Supplementary Figure 2).
- Drug dose data is captured using both an exponentially decaying sum, and a time since the last measurement. Decay rates with half-lives of 5 minutes and 60 minutes were used to capture both near term and longer average drug dosing effects.

To ensure that there was enough training data for each feature we removed features that had less than 100 recorded data values for the real-time model, and less than 50 for the initial model. For a full list of the 3,797 features used by Prescience for initial predictions see Supplementary Table 4 (Supp_InitialFeatureTable_4.csv). For the 3,905 features used in intraoperative predictions see Supplementary Table 5 (Supp_RealtimeFeatureTable_5.csv).

**Supplementary Fig 2.**
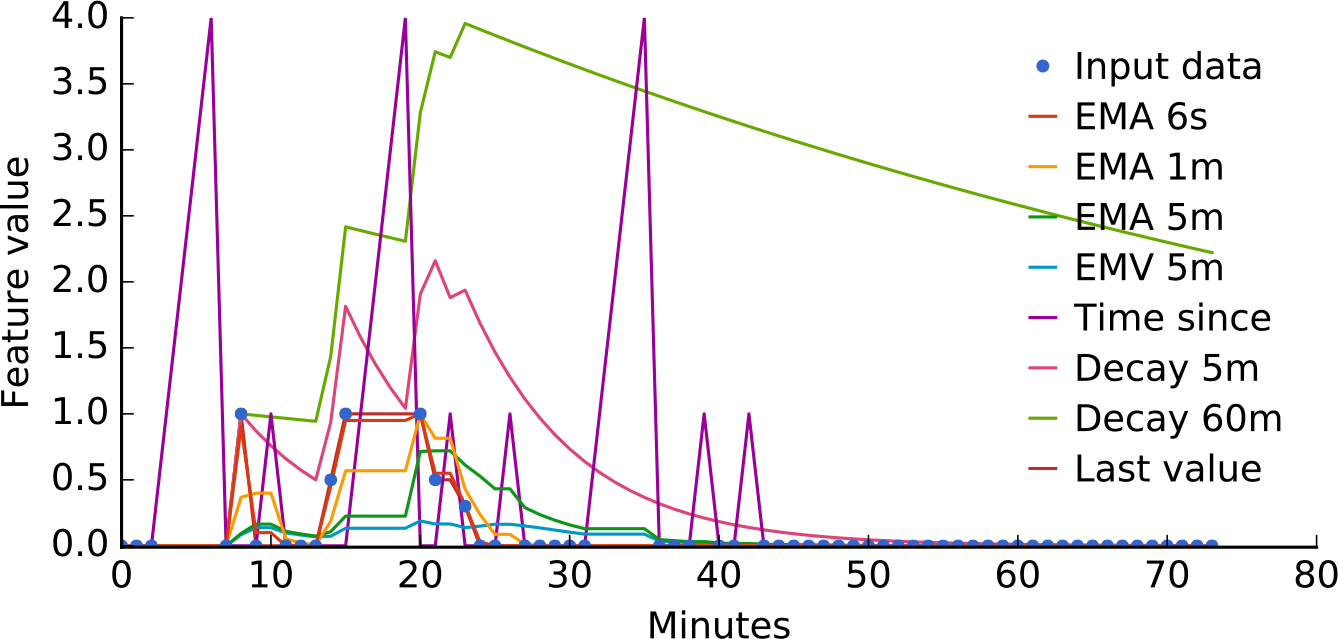
Responses of the eight different time series features used in Prescience to a sample set of unevenly reported data values. The blue dots represent the original unevenly sampled data, while curves represent the value of a feature over time. ‘EMA’ stands for exponential moving average and ‘EMV’ for exponential moving variance. Both EMA and EMV features are computed over weighted samples, where the weights decay with a specific half-life (6 seconds, 1 minute, or 5 minutes).

### Gradient boosting machines for prediction

The features we compute from real-time operating room data have a variety of complex nonlinear interactions. Capturing these requires a model with significant flexibility, and we chose a non-parametric approach called *gradient boosting machines (19*).

We compared the performance of gradient boosting against three baseline methods: Lasso penalized linear logistic regression; a linear SVM autoregressive model previously proposed for predicting hypoxemia based only on the SpO_2_ data stream (*9);* and an unsupervised Parzen window method used previously to predict patient deterioration (*17*). Gradient boosting machines significantly outperformed all baseline methods for our primary endpoint, real-time hypoxemia prediction (Supplementary Figure 3). For our secondary task of initial prediction gradient boosting machines were only slightly superior (Supplementary Figure 4). The large performance gain of gradient boosting for intraoperative prediction (Supplementary Figure 3) is likely because there are 8 million training samples, while for preoperative predictions (Supplementary Figure 4) there are only 42,000 samples and no time series data. Note that for initial prediction the autoregressive SVM and Parzen window methods were not applicable and hence not evaluated.

Gradient boosting machines are non-parametric models that draw a parallel between boosting and gradient descent in function space. They additively build up simpler models, like boosting, and these models are fit to the gradient of the loss at every data point. The most common type of basic model used is a regression tree because it is both robust to outliers and flexible. Taking some small fraction, *η*, of many trees fit to the gradient results in many small gradient descent steps in function space.

Fitting the trees is computationally challenging on large datasets so we used XGBoost, a recent high performance implementation of gradient boosting machines (*20*). For the real-time model we used *η* = 0.2 and 1,242 trees, while for the initial model we chose *η* = 0.1 and 4,000 trees. Using a smaller *η* value means more trees are required for fitting, which requires more time to run, but results in a smoother (and generally better) model. For both initial and real-time models we used bagging, where trees were trained on a random 50% subsample of the training data. For the preoperative model the max tree depth was 4 and the minimum child weight of any branch in the trees was 1. For the real-time model the max tree depth was 6 and the minimum child weight of any branch in the trees was 10.

All method parameters were tuned (and methods were chosen) using a validation set of operating room procedures separate from the final test set used for all final performance results. To ensure that there was no bias towards the final test set, the test data was initially compressed and left compressed until after method development was completed.

**Supplementary Fig 3.**
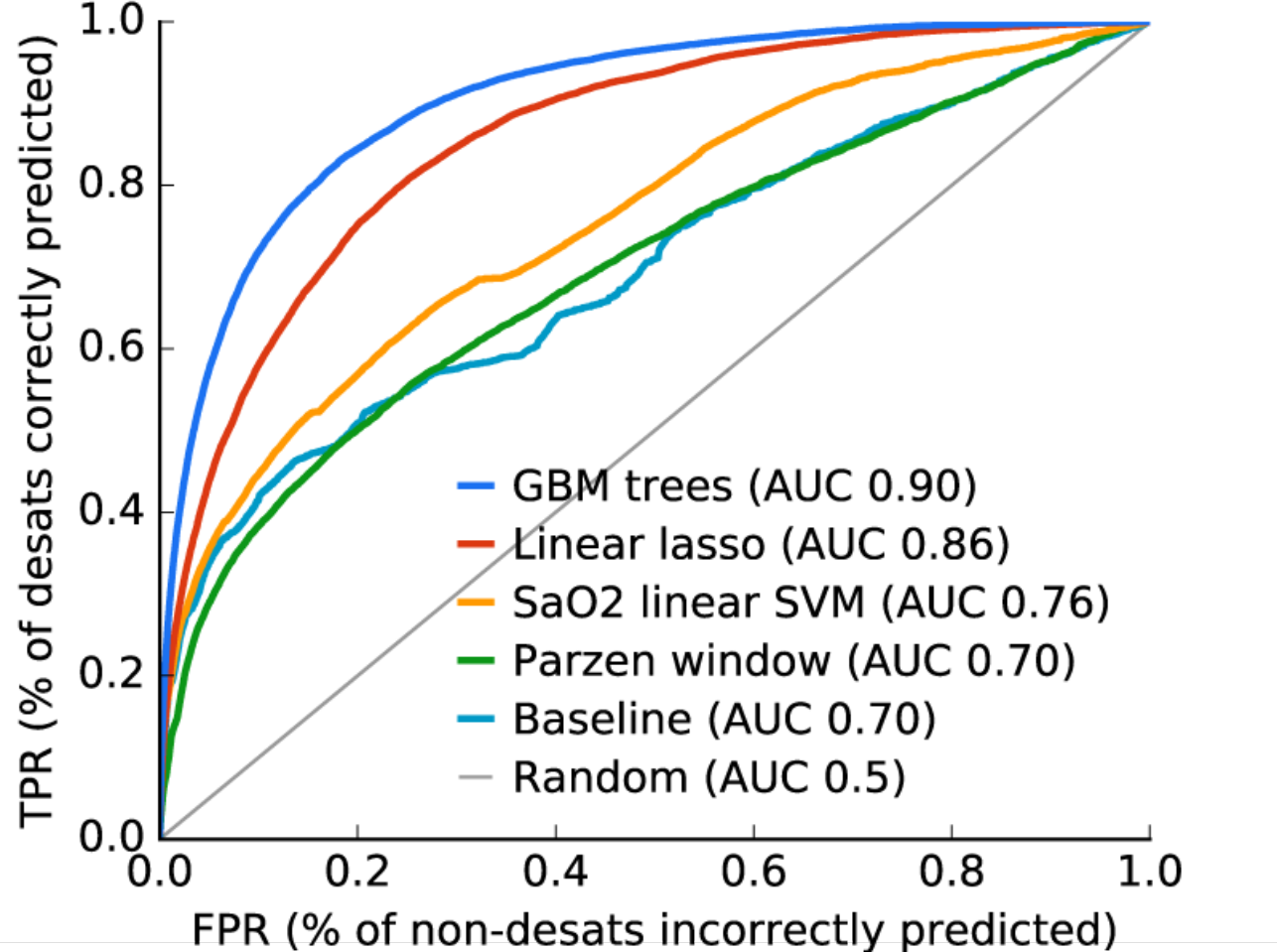
Real-time performance of gradient boosting machines vs a linear lasso model, a SVM based on ElMoaqet et al. (*9*), and an unsupervised Parzen window method used by Tarassenko et al. (*17*). There are ~8 million training samples and the increased flexibility of gradient boosting trees clearly outperforms the more restrictive linear models.

**Supplementary Fig 4.**
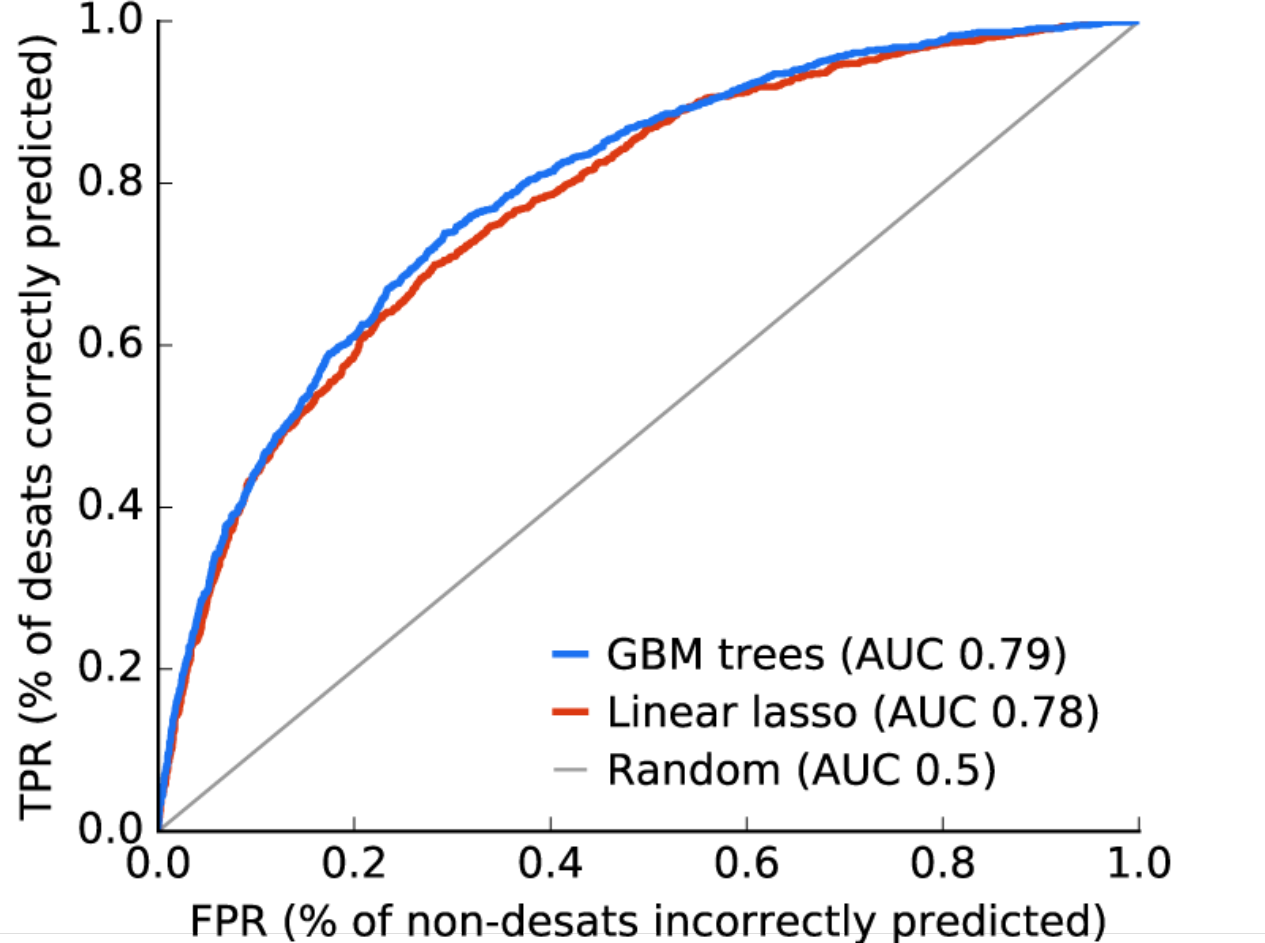
Preoperative performance of gradient boosting machines vs a linear lasso model. Given a much smaller preoperative dataset with ~42,000 training examples the difference between the complex gradient boosting trees model and a linear model becomes small.

### Computing feature importance estimates

Understanding why a statistical model has made a specific prediction is a key challenge in machine learning. It engenders appropriate trust in predictions and provides insight into how a model may be improved. However, many complex models with excellent accuracy, such as gradient boosting, make predictions even experts struggle to interpret. This forces a tradeoff between accuracy and interpretability. In response to this we chose to use a model agnostic representation of feature importance, where the impact of each feature on the model is represented using *Shapley values* (27), theoretically proven unique feature attribution values that have four important properties below. We developed an efficient method to compute the Shapely values (i.e., estimated importance of features to a particular prediction) in a real-time manner.

Shapley values are from the game theory literature and provide a theoretically justified method for allocation of credit among a group (see Eq (1)). In Prescience the group is a set of interpretable model input feature values, and the credit is the value of the prediction made by the model when given those input feature values. Feature impact is defined as the change in prediction probability when a feature is observed vs. unknown. Some feature values have a large impact on the prediction, while others have little impact. The Shapley values *ϕ*_*i*_(*f,x*), explaining a prediction *f*(*x*), are an allocation of credit among the various features (such as age, weight, time series features, etc.), and are the only such allocation that obeys a set of desirable properties (*14*, *16*). Given a prediction function *f(x*) we can define *f*_*x*_(*S*) = *f*(*x*_*s*_), where *x*_*s*_ is an input vector equal to *x* for features in the set *S* but equal to the missing data value otherwise.

Using this notation, we use the following set of properties:

**Efficiency.**

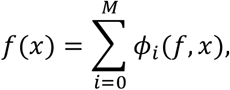

where *M* is the number of ‘interpretable’ inputs, which each correspond to a group of original inputfeatures. For Prescience these groups are the sets of features associated with each time series. For instance, the 6 second, 1 minute, and 5 minute moving average features, and the 5-minute moving variance feature from the SpO_2_ time series are all considered as a single group. The efficiency assumption forces the model to correctly capture the original predicted value.

**Symmetry.** If for all subsets *S* that don’t contain *i* or *j*

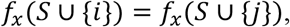

then *ϕ*_*i*_(*f*,*x*) = *ϕ*_*j*_(*f*,*x*). This states that if two features have an identical effect when observed in any situation then the Shapley values for the features must be the same.

**Null effects.** If for all subsets *S* that don’t contain *i*

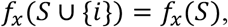

then *ϕ*_*i*_(*f*,*x*) = 0. This states that if a feature has no effect when observed in any situation then its Shapley value must be 0.

**Linearity.** For any two model prediction functions *f* and *f′*

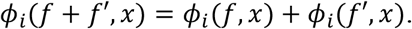

This states that the effect a feature has on the sum of two functions is the effect it has on one function plus the effect it has on the other.

Shapley showed that *only one allocation of credit satisfies these properties* and that allocation is the one given by the Shapley values.

Given a specific prediction *f*(*x*) we can compute the Shapley values using a weighted sum:

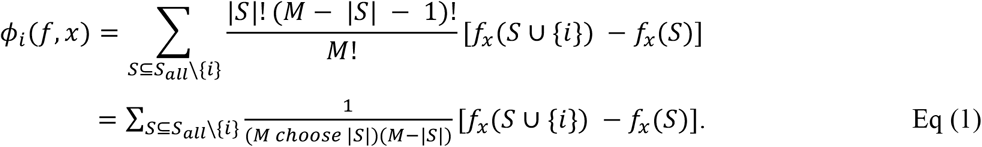

In practice, there are far too many terms to evaluate this sum completely, so we instead calculate it by only selecting the highest weight terms. This selection can be done greedily, stopping once the variance of the estimate becomes small enough.

To compute the Shapley values of each prediction we need to estimate the predictions of the model when specific input features are missing. Since the model was not trained to support missing values we approximate what the model would predict (if retrained on that subset of input features) by sampling from the training data set and replacing the missing features with the values they would have had in that sample. By averaging many such samples, we can estimate the expected value of *f*_*x*_(*S*) only using evaluations of *f*_*x*_(*S*_*all*_) where no features are missing.

The approach above requires nested sampling, once to estimate the Shapley value and then from each sample we again sample to estimate *f*_*x*_(*S*) and *f*_*x*_(*S* ∪ {*i*}). To reduce the number of samples in the inner step, we used k-medians to generate 20 medians of the entire dataset, and then performed a weighted evaluation for only these 20 summary inputs as an approximation for the entire dataset. This removes the need for nested sampling.

One final extension of Shapley value estimation we found to be helpful in Prescience was a non-linear link function *h* such that:

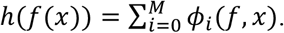

Since Prescience uses logistic regression the use of a *h* = *logit* link function transforms the output space from probabilities to log odds. Assuming the importance of features is additive in the log odds space is much more natural than assuming they are additive in the space of probabilities (which must fall between 0 and 1). The same reasoning also drives the use of the logit link function during standard logistic regression.

We were able to get stable feature importance estimates for thousands of features in less than 5 seconds on our server. We compared these theoretically grounded explanations with a simple estimate of feature importance to verify they showed reasonable consistency. The simple method we chose was to replace a single feature group with random values from other samples in the data set, and determine the average model output over different possible samplings. We then subtracted this mean value from the original model prediction to get a difference from a prediction with a typical value of that feature vs. the current value. This simple method is not very scalable and does not account for interactions with other features, but is useful to compare with the Prescience explanations to ensure the Prescience estimates of feature effects are consistent with an intuition of how much a feature’s change from its typical value effects the current risk of hypoxemia (Supplementary Figure 5).

**Supplementary Fig 5.**
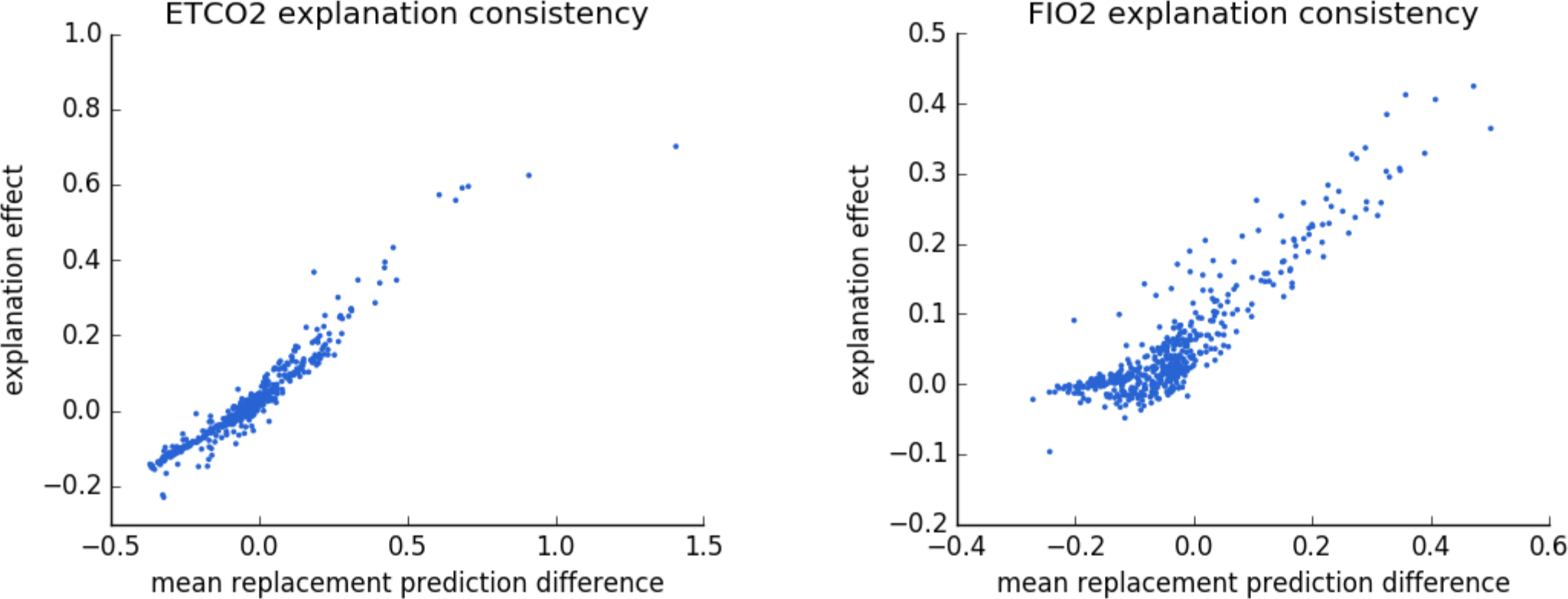
Consistency of Prescience explanation effects. Comparing the Prescience explanation effects with the difference between the current model output and the outputs when a specific feature is replaced with its typical value. The effects are from the cases shown to anesthesiologists during testing in Figure 2. The strong correlation (R^2^ = 0.92 for ETCO_2_ and R^2^ = 0.81 for FIO_2_) demonstrates the consistency of Prescience explanation effect sizes with the intuitive notion of the change in model out from replacing a feature’s value with a typical value. Note that both values are shown explaining the additive portion of the classification model (inside the logistic function).

### Physician evaluation

The potential benefit Prescience provides to physicians was evaluated using previously recorded procedures. Both before a procedure begins, and at several time points during the operation all the available electronically recorded data was shown to the anesthesiologist and they were asked to predict if a desaturation (as defined above) will occur in the next 5 minutes (Supplementary Figures 6, 7, and 8). For half of the procedures anesthesiologists are given Prescience explained risks, and for the other half they are given the same data, but without any Prescience assistance. In both cases anesthesiologists are asked to provide a fold change in the risk that desaturation will occur.

The test procedures were divided into two equal sized groups, replicate 1 and replicate 2. Anesthesiologists were also divided into two groups, A and B. Group A was given Prescience assistance on replicate 1 but not on replicate 2, while group B was given Prescience assistance on replicate 2 but not replicate 1. After randomly assigning anesthesiologists to groups, three anesthesiologists from group A completed the evaluation and two anesthesiologists from group B. We pooled the results within each group and between groups, and the results of this evaluation are shown in Figure 2. The order in which anesthesiologists were presented with cases was random across both replicate sets.

**Supplementary Fig 6.**
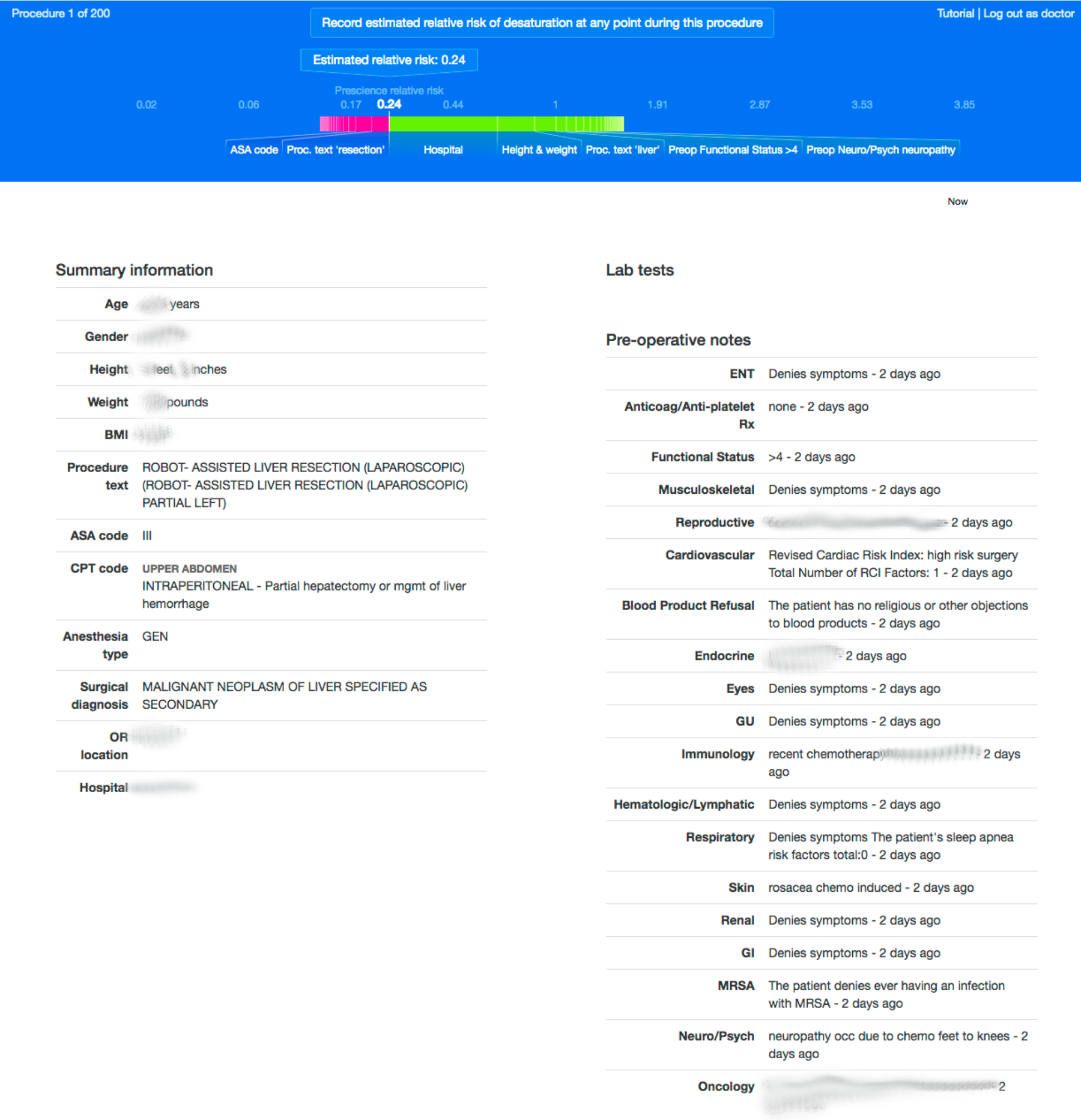
Sample of a physician’s test interface for preoperative prediction of risk. Prescience assistance is given for this preoperative prediction. Anesthesiologists choose an estimated relative risk by moving the given slider, then recording the score.

**Supplementary Fig 7.**
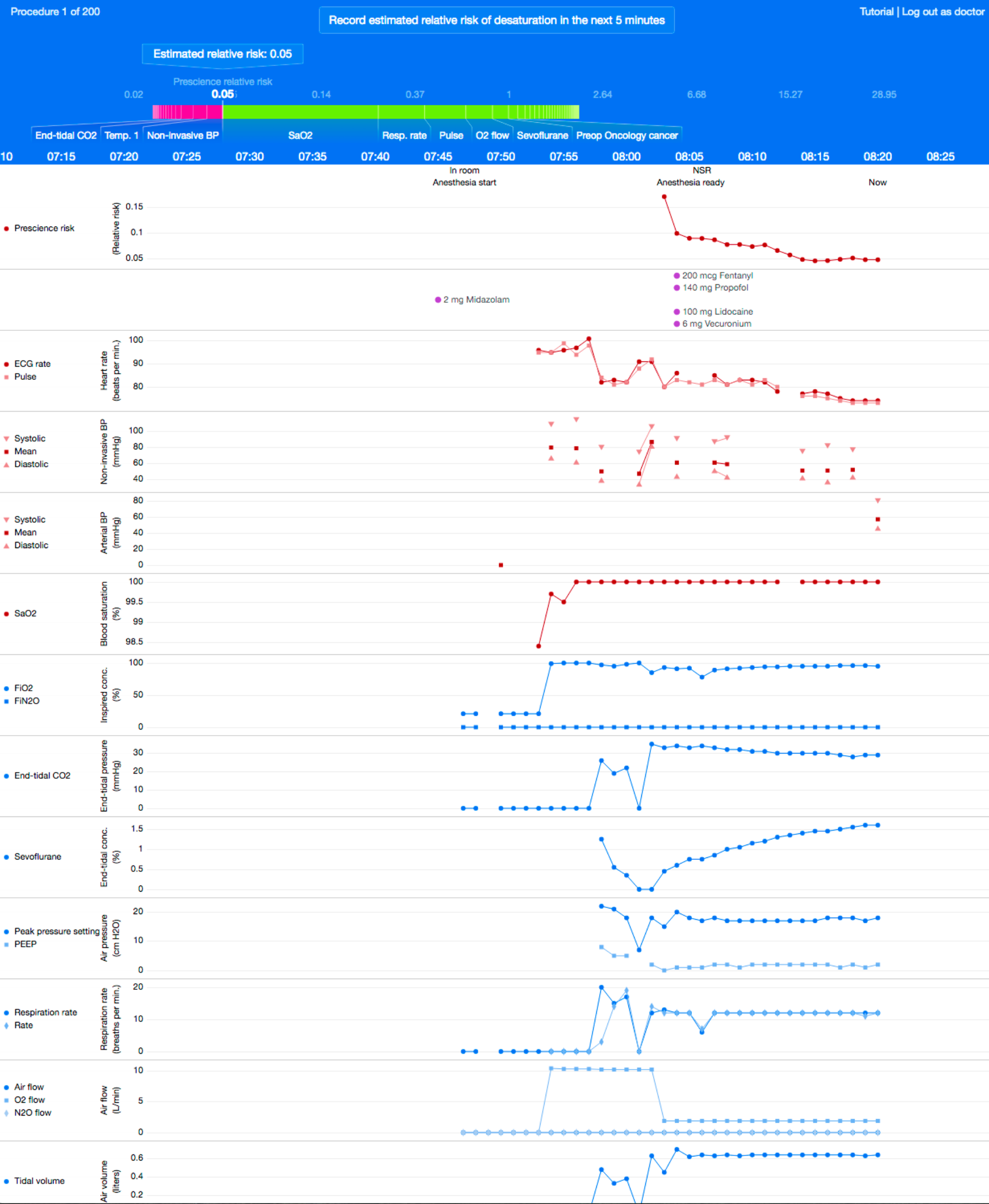
Sample of a physician’s test interface for real-time prediction of risk. Prescience assistance is given for this intraoperative prediction. Anesthesiologists choose an estimated relative risk by moving the given slider. The preoperative data is also shown lower down in the interface just as illustrated in Supplementary Figure 6.

**Supplementary Fig 8.**
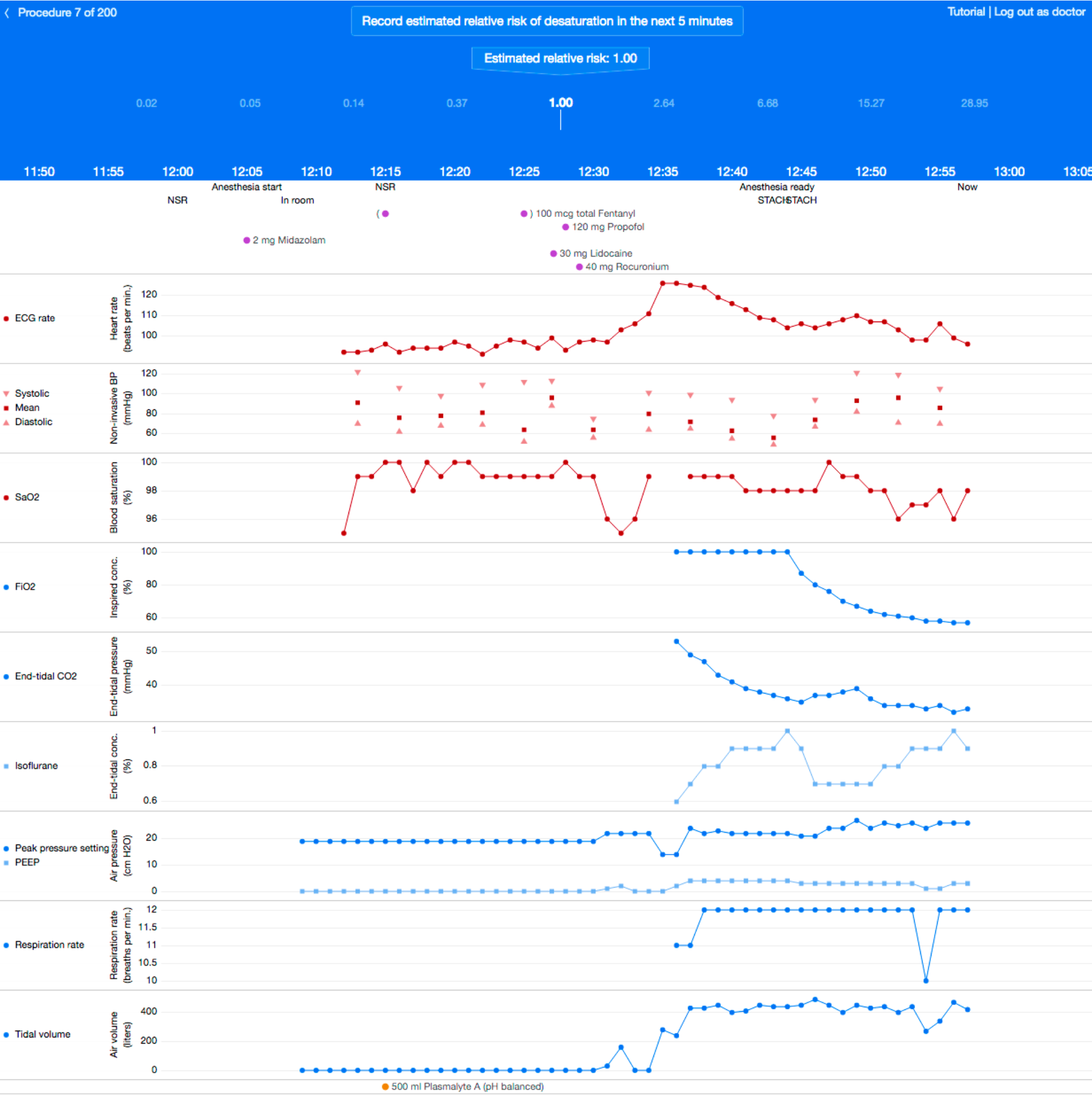
Sample of a physician’s test interface for real-time prediction of risk without Prescience assistance. This is similar to Supplementary Figure 7 but this time the anesthesiologist must choose a risk on their own using only the original data without help from Prescience.

**Supplementary Table 2.**
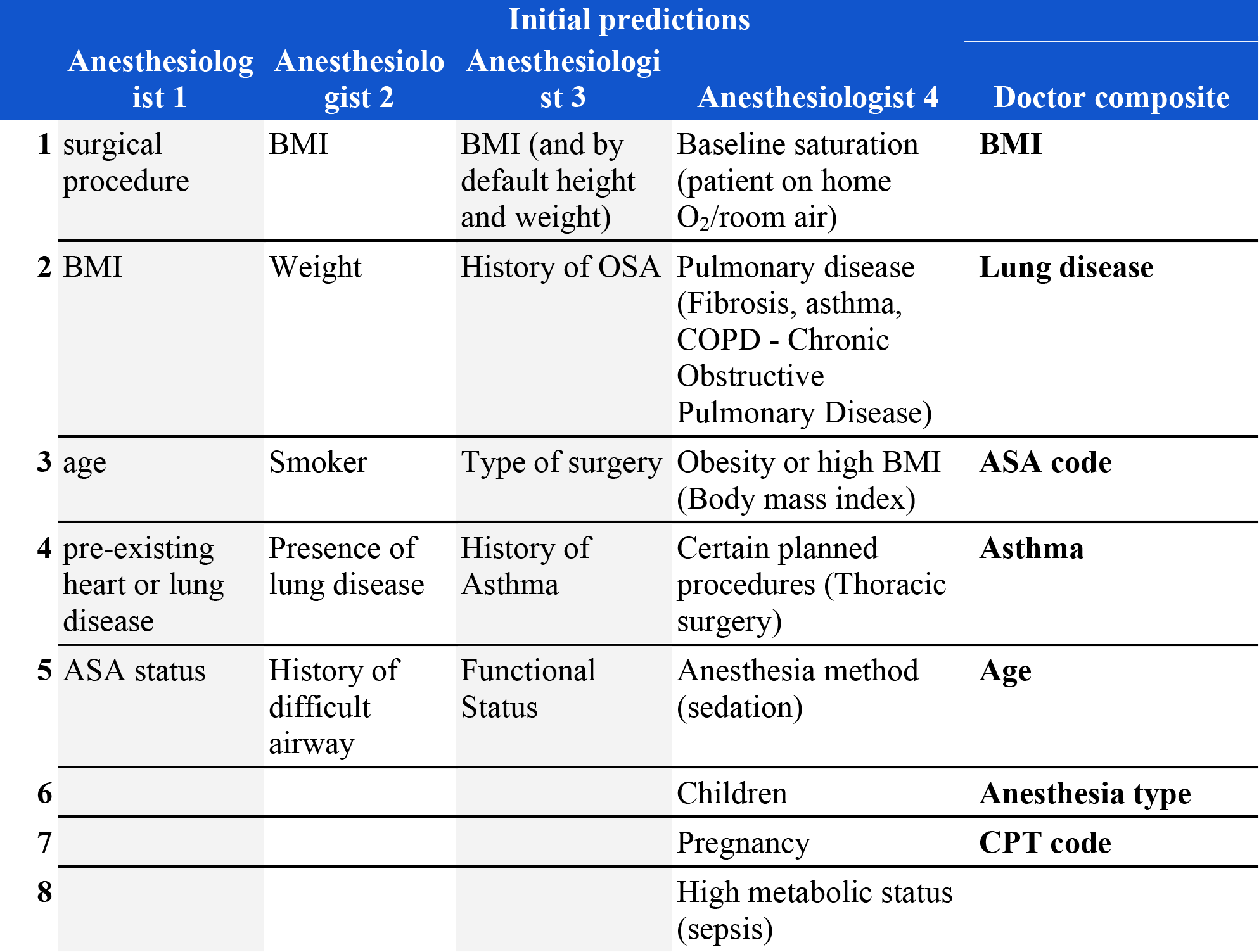
Physician’s responses listing the most important factors when assessing the risk for hypoxemia at any point during an upcoming procedure. Four anesthesiologists were asked to list the most important factors to consider for hypoxemia. The responses of all 4 anesthesiologists were then compiled into a single composite list. For a comparison with features chosen by Prescience, see Figure 4 in the main text.

**Supplementary Table 3.**
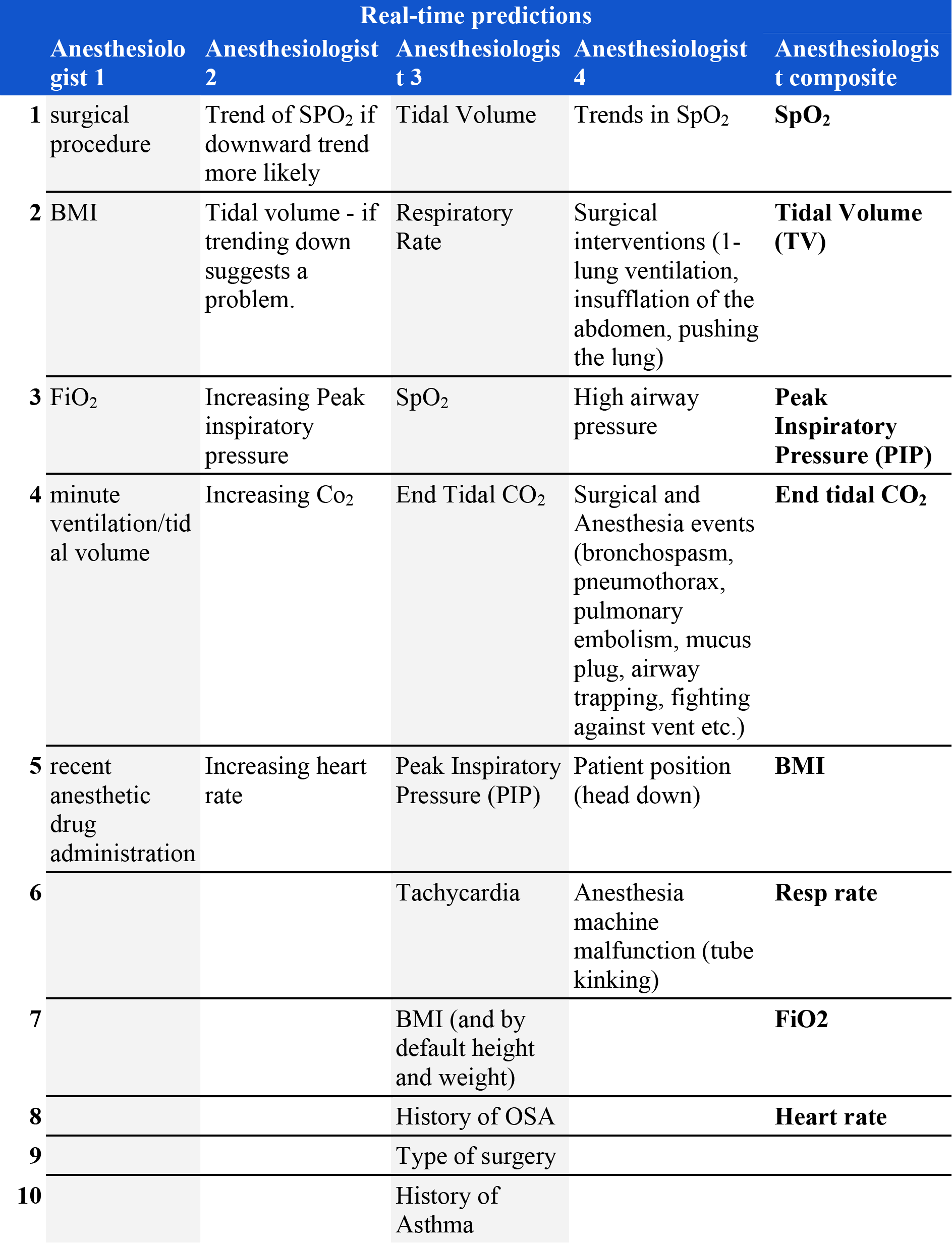
Physician’s responses listing the most important factors when assessing the risk for hypoxemia in the next 5 minutes. Four anesthesiologists were asked to list the most important factors to consider for hypoxemia. The responses of all 4 anesthesiologists were then compiled into a single composite list. For a comparison with features chosen by Prescience, see Figure 4 in the main text.

### Supplementary Materials

**Supplementary Fig 1. Criteria for defining testing labels.** When comparing the performance with anesthesiologists, only time points that clearly either desaturate or not are used. Hypoxemia involves dropping from ≥ 95% to ≤ 92% in the next 5 minutes. Not desaturating means remaining ≥ 95% for both the past 10 minutes and the next 10 minutes.

**Supplementary Fig 2. Responses of the eight different time series features used in Prescience to a sample set of unevenly reported data values.** The blue dots represent the original unevenly sampled data, while curves represent the value of a feature over time. ‘EMA’ stands for exponential moving average and ‘EMV’ for exponential moving variance. Both EMA and EMV features are computed over weighted samples, where the weights decay with a specific half-life (6 seconds, 1 minute, or 5 minutes).

**Supplementary Fig 3. Real-time performance of gradient boosting machines vs a linear lasso model, a SVM based on ElMoaqet et al. (*9*), and an unsupervised Parzen window method used by Tarassenko et al. (*17*).** There are ~8 million training samples and the increased flexibility of gradient boosting trees clearly outperforms the more restrictive linear models.

**Supplementary Fig 4. Initial performance of gradient boosting machines vs a linear lasso model.** Given a much smaller preoperative dataset with ~42,000 training examples the difference between the complex gradient boosting trees model and a linear model becomes small.

**Supplementary Fig 5. Consistency of Prescience explanation effects.** Comparing the Prescience explanation effects with the difference between the current model output and the outputs when a specific feature is replaced with its typical value. The effects are from the cases shown to anesthesiologists during testing in Figure 2. The strong correlation (R^2^ = 0.92 for ETCO_2_ and R^2^ = 0.81 for FIO_2_) demonstrates the consistency of Prescience explanation effect sizes with the intuitive notion of the change in model out from replacing a feature’s value with a typical value. Note that both values are shown explaining the additive portion of the classification model (inside the logistic function).

**Supplementary Fig 6. Sample of a physician’s test interface for initial prediction of risk.** Prescience assistance is given for this preoperative prediction. Anesthesiologists choose an estimated relative risk by moving the given slider, then recording the score.

**Supplementary Fig 7. Sample of a physician’s test interface for real-time prediction of risk.** Prescience assistance is given for this intraoperative prediction. Anesthesiologists choose an estimated relative risk by moving the given slider. The preoperative data is also shown lower down in the interface just as illustrated in Supplementary Figure 5.

**Supplementary Fig 8. Sample of a physician’s test interface for real-time prediction of risk without Prescience assistance.** This is similar to Supplementary Figure 6 but this time the anesthesiologist must choose a risk on their own using only the original data without help from Prescience.

**Supplementary Table 1. Raw data sources used for machine learning.** A wide variety of data sources were used that included text values, time series, discrete values, and quantitative values.

**Supplementary Table 2. Physician’s responses listing the most important factors when assessing the risk for hypoxemia at any point during an upcoming procedure.** Four anesthesiologists were asked to list the most important factors to consider for hypoxemia. The responses of all 4 anesthesiologists were then compiled into a single composite list. For a comparison with features chosen by Prescience, see Figure 4 in the main text.

**Supplementary Table 3. Physician’s responses listing the most important factors when assessing the risk for hypoxemia in the next 5 minutes.** Four anesthesiologists were asked to list the most important factors to consider for hypoxemia. The responses of all 4 anesthesiologists were then compiled into a single composite list. For a comparison with features chosen by Prescience, see Figure 4 in the main text.

**Supplementary Table 4. Preoperative features used by Prescience.** An enumeration of all the 3,797 features used for preoperative predictions (Supp_InitialFeatureTable_4.csv).

**Supplementary Table 5. Intraoperative features used by Prescience.** An enumeration of all the 3,905 features used for intraoperative predictions (Supp_RealtimeFeatureTable_5.csv).

## Code availability

All modeling and processing code is available from the authors upon request. However, note that patient privacy prevents training data from accompanying the code.

## Data availability

The operating room datasets from participating hospitals are not publically available due to patient privacy concerns. For more information on obtaining additional summaries of the data please contact the corresponding author.

## Footnotes

1 In Figure 4A the 2nd and 4th features chosen by anesthesiologists (lung disease and asthma, respectively) were not well captured in the AIMS data, and did not show up as important in the Prescience model.

